# Development and extensive sequencing of a broadly-consented Genome in a Bottle matched tumor-normal pair

**DOI:** 10.1101/2024.09.18.613544

**Authors:** Jennifer H. McDaniel, Vaidehi Patel, Nathan D. Olson, Hua-Jun He, Zhiyong He, Kenneth D. Cole, Alexander A. Gooden, Anthony Schmitt, Kristin Sikkink, Fritz J Sedlazeck, Harsha Doddapaneni, Shalini N. Jhangiani, Donna M. Muzny, Marie-Claude Gingras, Heer Mehta, Sairam Behera, Luis F Paulin, Alex R Hastie, Hung-Chun Yu, Victor Weigman, Alison Rojas, Katie Kennedy, Jamie Remington, Isai Salas-González, Mitch Sudkamp, Kelly Wiseman, Bryan R. Lajoie, Shawn Levy, Miten Jain, Stuart Akeson, Giuseppe Narzisi, Zoe Steinsnyder, Catherine Reeves, Jennifer Shelton, Sarah B. Kingan, Christine Lambert, Primo Bayabyan, Aaron M. Wenger, Ian J. McLaughlin, Aaron Adamson, Christopher Kingsley, Melanie Wescott, Young Kim, Benedict Paten, Jimin Park, Ivo Violich, Karen H Miga, Joshua Gardner, Brandy McNulty, Gail L. Rosen, Rajiv McCoy, Francesco Brundu, Erfan Sayyari, Konrad Scheffler, Sean Truong, Severine Catreux, Lesley Chapman Hannah, Doron Lipson, Hila Benjamin, Nika Iremadze, Ilya Soifer, Gat Krieger, Stephen Eacker, Mary Wood, Erin Cross, Greg Husar, Stephen Gross, Michael Vernich, Mikhail Kolmogorov, Tanveer Ahmad, Ayse Keskus, Asher Bryant, Francoise Thibaud-Nissen, Jonathan Trow, Jacqueline Proszynski, Jeremy Wain Hirschberg, Krista Ryon, Christopher E. Mason, Mital S. Bhakta, J. Zachary Sanborn, Elizabeth M. Munding, Justin Wagner, Chunlin Xiao, Andrew S. Liss, Justin M. Zook

## Abstract

The Genome in a Bottle Consortium (GIAB), hosted by the National Institute of Standards and Technology (NIST), is developing new matched tumor-normal samples, the first to be explicitly consented for public dissemination of genomic data and cell lines. Here, we describe a comprehensive genomic dataset from the first individual, HG008, including DNA from an adherent, epithelial-like pancreatic ductal adenocarcinoma (PDAC) tumor cell line and matched normal cells from duodenal and pancreatic tissues. Data for the tumor-normal matched samples comes from seventeen distinct state-of-the-art whole genome measurement technologies, including high depth short and long-read bulk whole genome sequencing (WGS), single cell WGS, and Hi-C, and karyotyping. In future publications, these data will be used by the GIAB Consortium to develop matched tumor-normal benchmarks for somatic variant detection. We expect these data to facilitate innovation for whole genome measurement technologies, *de novo* assembly of tumor and normal genomes, and bioinformatic tools to identify small and structural somatic mutations. This first-of-its-kind broadly consented open-access resource will facilitate further understanding of sequencing methods used for cancer biology.

## Background & Summary

The GIAB Consortium has previously developed extensive sequencing data and benchmarks for seven normal human cell lines, which are used in technology development and validation of clinical assays.^1,2^ While some tumor cell lines have been characterized as benchmarks by other groups,^3–6^ these data are from legacy cell lines with no consent or consent before whole genome sequencing was routine. In 2013, the NIH determined that researchers could publicly share genomic data from legacy cancer cell lines in the Cancer Cell Line Encyclopedia despite having no or limited consent, because most of these cell lines already had public genomic data available (https://www.cancer.gov/ccg/research/structural-genomics/tcga/history/policies/ccle-open-release-justification.pdf). However, the policy states that it could be revised in the future, and it “strongly recommended that informed consent processes for prospective research development of human cell lines (for research or commercial purposes) fully describe and consider any risks associated with broad distribution of genomic data derived from those cell lines.” Therefore, there is a need for new tumor-normal pairs with explicit consent for public dissemination of genome sequencing data and cell lines, so that they can be developed into enduring reference samples for benchmarking somatic variants.^7^

To meet this need, we describe a comprehensive dataset of whole genome-scale measurements, including extensive short and long-read sequencing, from a new immortal PDAC cell line and paired normal pancreatic and duodenal tissues, derived from a female of European genetic ancestry with explicit consent for public release of genomic data. We will make additional data and benchmarks available as they are generated. We are also actively working to deposit the cell line in a public repository. Information and links for these resources as they become available, will be posted on the NIST Cancer Genome in a Bottle webpage (https://www.nist.gov/programs-projects/cancer-genome-bottle).

## Methods

### Pancreatic cancer patient with explicit consent for genomic data sharing

The GIAB Consortium requires explicit consent for genomic data sharing for its widely-used cell lines, so NIST is collaborating with the Liss Laboratory at Massachusetts General Hospital (MGH) (https://www.massgeneral.org/surgery/gastrointestinal-and-oncologic-surgery/research/liss-laboratory) to develop matched tumor-normal cell lines. The Liss Lab operates the Pancreatic Tumor Bank at MGH. MGH recruited participants, performed tissue collection and attempted to develop matched tumor-normal cell lines from the resected tissues (though normal cell line development was unsuccessful for the participant (GIAB ID: HG008) from which data is presented in this manuscript). The recruitment process for this study involved identifying adult patients with pancreatic disease treated at the Massachusetts General Hospital. Patients were approached by their treating surgeon or research nurse during their initial or preoperative office visits, at which time they were informed about the study. The surgeon or research nurse emphasized that participation was voluntary and would not affect the patient’s care at the hospital. The study was conducted in accordance with the Declaration of Helsinki. The participants have provided informed consent explicitly for tissue collection and public genomic data sharing, understanding the risks associated with these. Collection of tissue and blood from patients with pancreatic disease was approved by the Mass Brigham General IRB under protocol # 2003P001289. The MGH Data and Tissue Sharing Committee and NIST Research Protections Office determined that the signed consent allows for sharing of the participants’ genomic data and development and distribution of cell lines as reference materials. Relevant excerpts from the patient consent are provided below.

### Consent excerpt for sharing genomic data and cell line

For transparency, we include the parts of the MGH IRB-approved consent relevant to public sharing of genomic data, creation of the immortalized cell line, and public sharing of the cell line below:

“We plan to do genetic research on the DNA in your tissue sample. DNA is the material that makes up your genes. All living things are made of cells. Genes are the part of cells that contain the instructions which tell our bodies how to grow and work, and determine physical characteristics such as hair and eye color. Genes are passed from parent to child.”

“Your tissue sample may be used to create living tissue samples (including cell lines) that can be grown in the laboratory. This allows researchers to have an unlimited supply of your cells in the future without asking for more samples from you. The living tissue samples will be shared with the academic, non-profit research, and for-profit communities. Cancer researchers will use these living tissue samples to study better ways to detect and treat cancer. Companies may use the living tissue samples to develop products that are used to improve the detection and treatment of cancer to benefit patients. The living tissue samples will be sent with the random number corresponding to your original tissue and will not be sent with identifiable health information.”

“Your cells might be used in research involving genetic alteration of the cells. Your cells might be mixed with other human cells, mixed with animal cells, or grown in lab animals like mice. We may also perform a whole genome analysis on the DNA from your clinical samples and living tissue samples.

Usually researchers study just a few areas of your genetic code that are linked to a disease or condition. In whole genome studies, all or most of your genes are analyzed and used by researchers to study links to many diseases or conditions.”

“In order to allow researchers to share test results, the National Institutes of Health (NIH) and other central repositories have developed data (information) banks that analyze data and collect the results of whole genome studies.

These banks may also analyze and store DNA samples, as well. These central banks will store your genetic information and samples and make them available to the public for additional studies. Because the DNA sequence is available on a database, it is possible that someone using the sequence could identify the donor and possible relatives. The samples and data will be sent with only your code number attached. Your name or other directly identifiable information will not be given to central banks. There are many safeguards in place to protect your information and samples while they are stored in repositories and used for research.”

“Research using your samples, living tissue samples, and their whole genome information is important for the study of virtually all diseases and conditions. Therefore, the sample/data banks will provide study data for researchers working on any disease.”

### HG008 patient history

This dataset is for the first tumor cell line developed from an MGH participant (GIAB ID: HG008). The individual, known by the GIAB as HG008, was a 61-year-old female with XX sex chromosomes who was diagnosed with pancreatic ductal adenocarcinoma (disease stage ypT2 N1). After neoadjuvant therapy (gemcitabine and cisplatin followed by a single round capecitabine concurrent with radiation therapy), the greatest dimension of the tumor was 3.2 cm, located in the pancreatic head near the duodenum. The histologic grade was G2 (moderately differentiated), and the tumor regression grade was 2 (Minimal response: Residual cancer outgrown by fibrosis).

Resection of the PDAC tumor, duodenal tissue, and pancreatic tissue after neoadjuvant therapy was performed at MGH in December 2020 and sent to the Liss laboratory for tissue banking. There was one positive lymph node out of 13 examined. Perineural and large vessel (venous) invasion were present, confirmed on multiple elastic stains, but small vessel (blood/lymphatic) invasion was absent. Immunohistochemical stains for DNA mismatch repair proteins show preserved expression of all proteins (hMLH1, hMSH2, hMSH6, and hPMS2).

### Establishing the HG008-T tumor cell line

The Liss laboratory established a PDAC tumor cell line (HG008-T) from the resected PDAC tumor after neoadjuvant therapy. PDAC tumors are stromal-rich, with cancer cells often only composing 10 % of the tumor.

Approximately 50 mg of tumor tissue was enzymatically dissociated to begin culturing from the primary tumor cells for cell line development. While PDAC cell cultures often contain substantial fibroblasts, in this case all cells appeared to be epithelial-like tumor cells even at passage 2 for this cell line. Cells were cultured for the first 20 passages using “Rich” media composed of DMEM/F12 with 20 % Fetal Bovine Serum (FBS), Nicotinamide (0.01 M), cholera toxin (8.4 ng/mL), Epidermal Growth Factor (EGF; 10 ng/mL), Hepatocyte Growth Factor (HGF: 25 ng/mL), human recombinant insulin (10 ug/mL), human recombinant transferrin (5.5 ug/mL), and selenious acid (6.7 ng/mL). Cells were enzymatically removed from the culture surface using 0.25% trypsin and split by diluting no more than 1:3 for the first 20 passages. Passages 21-24 were diluted 1:3 and grown in the same media as prior passages however without EGF. Morphologically pure cancer cell cultures, free of tumor-associated fibroblasts, were considered cell lines after growing for at least 20 passages. From passage 25 on, the established cell line was then cultured in DMEM/F12 with 10 % FBS. All media contained 1 % antibiotic/antimycotic.

Once the cell line was established, a large homogenous batch (batch 0823p23) of tumor cells was cultured and at passage 23, approximately 80 million tumor cells were harvested, pelleted and snap-frozen for distribution and measurement of non-viable cells. Tumor cell line pellets, or isolated DNA from the pellets, and non-viable paired normal tissue, or isolated DNA from the tissue, were distributed for genome-scale measurements as described in Table 2 and Table S1 (see Supplementary Information document). In addition to measurements from the large, homogeneous batch of tumor cells, we also performed karyotyping on p31, Illumina WGS on p36, and Phase Genomics Hi-C on p38, all derived from the same series of passages starting with the p13 stock that batch 0823p23 was grown from. Passage 13 of the tumor cell line was transferred to NIST in summer of 2024 and preliminary measurements of NIST passages p18 and p21 are also presented (Figure 1).

**Figure 1.**
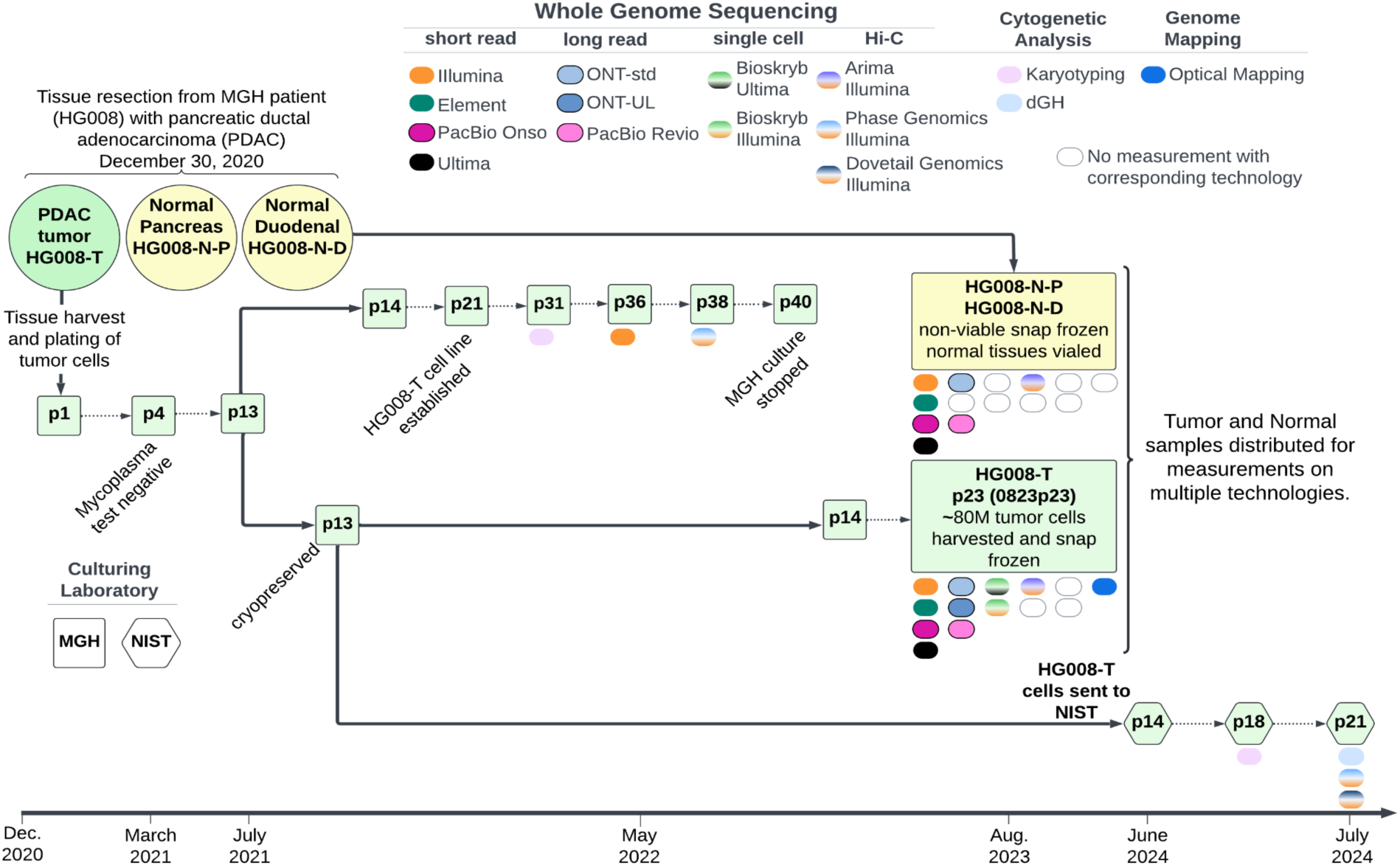
HG008 Passaging and Measurements. The PDAC tumor and normal duodenal and pancreatic tissues were resected from the HG008 individual in 2020. The HG008-T cell line was established by MGH and a bulk growth of tumor cells was produced for most measurements. This bulk growth and harvest is known as batch 0823p23. Tumor and normal samples were sent for measurements on multiple technologies denoted by the colored ovals; empty ovals are where no corresponding measurement was made. To characterize the HG008-T cell line during passaging, preliminary measurements were made and noted below the passage points denoted by “p#”. In 2024, an aliquot of the cell line was transferred to NIST by MGH for additional culturing and measurements.

### HG008 paired normal tissues

For the HG008 participant, the laboratory attempted to create an EBV-immortalized lymphoblastoid cell line from frozen PBMCs, but this was unsuccessful. However, several hundred milligrams of normal pancreatic (HG008-N-P) and duodenal (HG008-N-D) tissue was resected, allowing for matched normal sample measurements. These normal sample measurements are primarily to help identify somatic variants in the tumor, but also could be useful for mosaic variant detection within the tissues.

### Tumor and normal samples for measurement

Tumor and normal samples required genomic DNA (gDNA) isolation prior to measurement. In some cases measurement sites performed their own DNA isolation, while others received isolated gDNA. The samples the measurement sites received, either DNA, cells or tissue, are noted in Table 2. For sites receiving gDNA, isolations from non-viable cell pellets of the tumor cell line (batch 0823p23), non-viable normal duodenal tissue and non-viable pancreatic tissue were performed by the Genome Technology laboratory at Northeastern University and distributed to the measurement sites. Methods for measurement sites that performed their own DNA isolations are provided with their measurement methods in the Genome-scale measurements methods section. For sites receiving gDNA, methods for normal tissue and tumor cell line gDNA isolations are provided below and referenced in Genome-scale measurement methods as “previously isolated genomic DNA.”

### Genomic DNA isolation from normal tissues

Genomic DNA (gDNA) was isolated from two normal tissue types, non-viable pancreatic (HG008-N-P) and duodenal (HG008-N-D), using the Qiagen Puregene Tissue Kit (PN: 158667). Approximately 2 mL of Qiagen kit cell lysis buffer with 30 uL of Proteinase K was added to 10-15 mg of tissue. A Dounce Homogenizer was used to disrupt the tissue before leaving it overnight at 55℃. This was followed by addition of RNase A and another incubation at 37℃ for 5 minutes followed by cooling on ice. Genomic DNA was precipitated, washed and hydrated in TE buffer (1 mM EDTA, 10 mM Tris·Cl pH 7.5) per the Qiagen Puregene protocol. Two separate isolations were performed for each tissue type and resulting gDNA pooled for each tissue type. For QC, the isolating laboratory measured the yield of gDNA by Qubit and read N50 of gDNA with Oxford Nanopore Technologies (ONT) sequencing. Further, sites performing long-read sequencing independently assessed the gDNA prior to use. Yield, read N50 and sizing are summarized in Table 1. Isolated gDNA from the paired normal tissues was then distributed to various sites for genome-scale measurements as described in Table 2.

**Table 1.** Batch genomic DNA (gDNA) isolation for distribution. DNA isolations from the three HG008 samples were performed for distribution to laboratories for measurement. Isolated DNA was sent to several sites for genome-scale measurements as described in Table 2. To confirm gDNA was fit for purpose, yield and length of gDNA was measured at different sites prior to use. Sizing and N50 reported are approximated values from three measurement sites. Genomic DNA yield from normal tissues for the separate isolations is noted in parentheses.

| Sample Input | Measurement Method | HG008-T | HG008-N-D | HG008-N-P |
| --- | --- | --- | --- | --- |
|  |  | 5 million non-viable tumor cells from batch 0823p23 | two separate isolations of 10 mg of normal duodenal tissue were performed | two separate isolations of 10 mg of normal pancreatic tissue were performed |
| Genomic DNA Yield | Qubit | 60 µg | 34 µg<br>(18 µg and 16 µg) | 16 µg<br>(5 µg and 11 µg) |
| Read N50 | ONT sequencing | 30 kb | 10 kb | 8 kb |
| Sizing | FemtoPulse | 58 kb | NA | 26 kb |
|  | Tapestation | NA | 53 kb | NA |

**Table 2.**
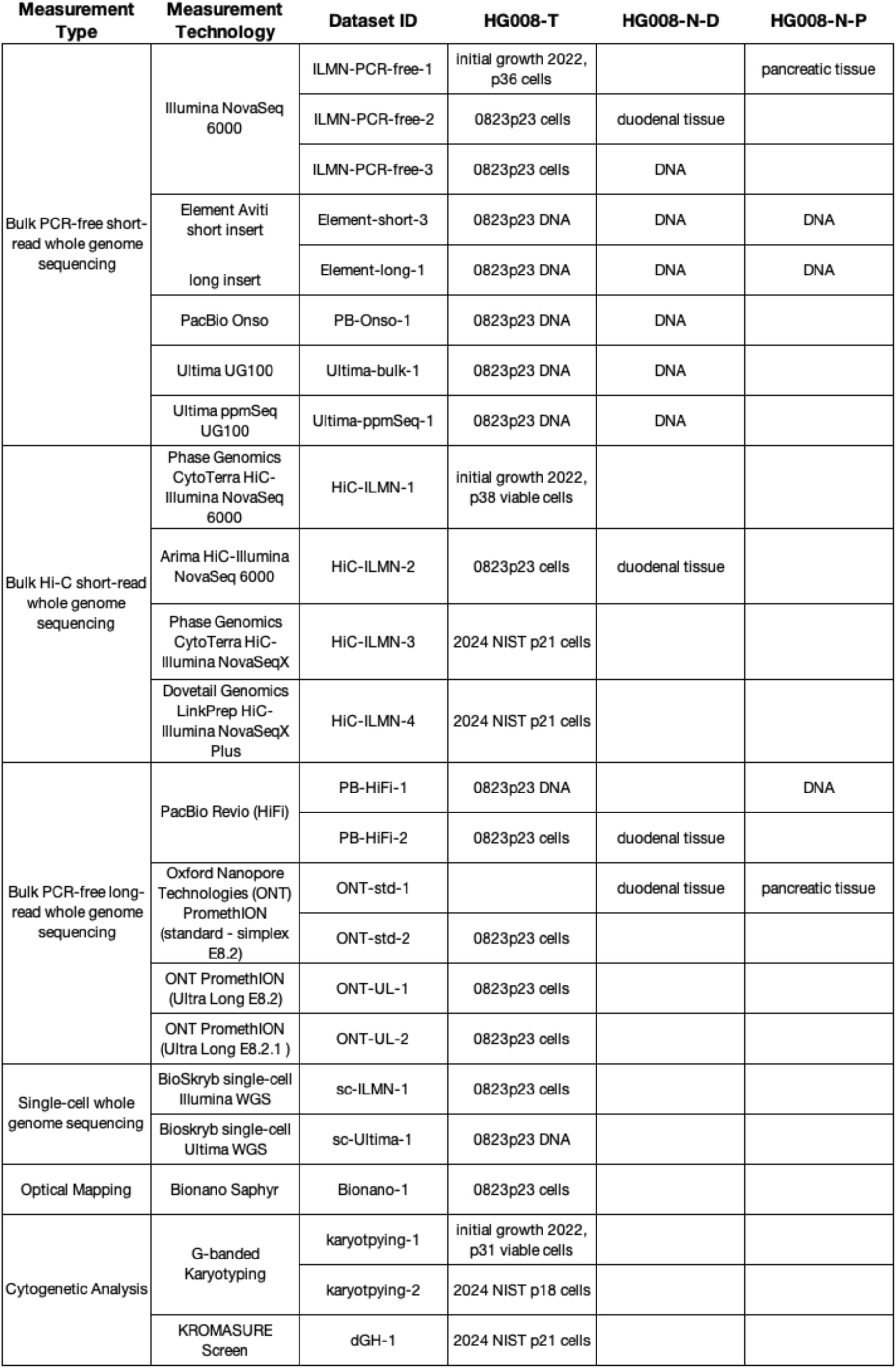
Summary of measurements made for HG008 tumor and normal samples. Sample fields denote the starting material received by the measurement institution; cells, tissue or isolated genomic DNA. Most tumor samples derive from a large, homogeneous batch of cells distributed in 2023 (0823p23) by the Liss lab at MGH. However, some are from a series of Liss lab passages in 2022 or from NIST passages in 2024, which were both started from the same Liss lab p13 cryopreserved cells used to create batch 0823p23, as shown in Figure 1.

### Genomic DNA isolation from the tumor cell line

Genomic DNA (gDNA) was isolated from non-viable cell pellets of the 0823p23 batch of cells from the HG008-T tumor cell line using the Qiagen Puregene Cell Kit (PN: 158767). Approximately five million cells were input into the Qiagen Puregene protocol. Briefly, cells were lysed, treated with RNase, gDNA precipitated, washed and hydrated in TE buffer (1 mM EDTA, 10 mM Tris·Cl pH 7.5). The gDNA was viscous and was therefore allowed to homogenize for one week at ambient laboratory temperature. Yield of gDNA was measured by Qubit. N50 of gDNA isolated from the tissues was estimated with ONT sequencing (Table 1). Isolated genomic DNA from the 0823p23 batch of HG008-T cells was then distributed to various sites for genome-scale measurements as described in Table 2.

### Genome-scale measurements of HG008 tumor and normal samples

Extensive genomic measurements were performed for all HG008 samples; HG008-T, HG008-N-D and HG008-N-P (Figure 1). Here we present 24 GIAB HG008 whole genome-scale datasets. Measurements were performed with 14 collaborating laboratories representing 17 distinct measurement technologies (summary in Table 2 and full description in Table S1, see Supplementary Information document). The measurement methods presented below include samples measured (HG008-T, HG008-N-P and/or HG008-N-D), DNA isolation, sample preparation, measurement process and validation, as appropriate for the measurement.

### Bulk short-read whole genome sequencing^8^

Illumina NovaSeq 6000

### PCR-free 2×150 bp Illumina Sequencing (Dataset ID: ILMN-PCR-free-1)

#### DNA Isolation

Genomic DNA (gDNA) was isolated from approximately 10 million non-viable HG008-T cells (passage 36 from 2022 culture) and 80 mg of non-viable HG008-N-P tissue with the Qiagen Gentra Puregene kit using the manufacturer’s instruction. Genomic DNA mass yields for tumor and normal samples were approximately 26 and 9 ug, respectively. Resulting DNA was quantified using the PicoGreen Assay (Quant-iT™ PicoGreen™, catalog# P11496) and the quality was assessed using 1 % agarose gel (Invitrogen™ E-Gel™ Agarose Gels, catalog# G720801).

#### Library Preparation

Library preparation was performed for each sample using an in-house PCR-free protocol, with the Roche Kapa HyperPrep PCR-free Kit (PN: KK8505). Two libraries were prepared for the tumor from 1 ug of gDNA for each library and one library was prepared for the normal duodenal sample from 1 ug of gDNA. For both tumor and normal samples, DNA was sheared into fragments of approximately 450 to 600 bp using a Covaris E220 system.

Fragmentation was followed by purification of the fragmented DNA using double-sided bead clean up with different ratios of AMPure XP beads to remove very small and large fragments. DNA end-repair and 3’-adenylation were performed in the same reaction followed by ligation of the barcoded adapters using a set of 96 8-bp barcoded adapters from IDT, Illumina TruSeq RNA unique dual (UD) Indexes (PN: 20040870). Post ligation products were cleaned twice with AMPure XP beads to get rid of leftover adapters/adapter dimers.

#### Whole Genome Sequencing

The resulting tumor and normal PCR-free libraries were sequenced together in one lane using the Illumina NovaSeq 6000 instrument (ICS v1.8.1 and bcl2fastq2 version 2.20.0.422).^9^ Paired-end, 2×150 bp sequencing, was performed using the Illumina NovaSeq 6000 S4 Reagent Kit v1.5 - 300 cycles (PN: 20028312) with the NovaSeq XP 4-Lane Kit v1.5 (PN: 20043131).

#### Validation Methods

The Illumina sequencing paired-end FASTQs were assessed with FastQC. The Illumina FASTQs were aligned to the GRCh38 reference genome using BWA-MEM. The GA4GH QC pipeline was run on the resulting BAM files to assess the quality metrics of the alignments (Table S2, see Supplementary Information document). A summary of QC metrics can be found Table 3.

**Table 3.** Short-read metrics. Summary of metrics generated by the GA4GH Quality Control pipeline, Samtools and Mosdepth for validation of short-read PCR-free data. The Samtools error rate measures substitution and indel differences between the reads and the reference, including differences resulting from true variants. All mapping-based metrics presented are based on alignments to the GRCh38 reference genome. *The Onso percent properly paired and percent mapped reads metrics presented represents pairing metrics following filtering of 1 bp reads described with the Onso methods. Onso data presented as part of the HG008 dataset is not filtered. **Element-short-1 and Element-short-2 datasets are superseded by Element-short-3, which utilizes a commercially available sequencing chemistry and addresses a previous reagent issue.

| Technology | Dataset ID | GIAB sample ID | samtools error rate (%) | average length (bp) | avg. insert size (bp) | insert size SD | properly paired (%) | mapped reads (%) | mean autosome coverage | diploid mean coverage | haploid mean coverage | NRPCC |
| --- | --- | --- | --- | --- | --- | --- | --- | --- | --- | --- | --- | --- |
| Illumina NovaSeq 6000 WGS PCR-free 2x150 | ILMN-PCR-free-1 | HG008-N-P | 0.6 | 151 | 444 | 102 | 97.2 | 99.6 | 150 |  |  |  |
|  |  | HG008-T | 0.5 | 151 | 427 | 97 | 97.6 | 99.8 | 140 | 191 | 96 | 48 |
|  | ILMN-PCR-free-2 | HG008-N-D | 0.5 | 151 | 417 | 94 | 96.6 | 99.7 | 169 |  |  |  |
|  |  | HG008-T | 0.6 | 151 | 432 | 97 | 96.8 | 99.7 | 141 | 195 | 98 | 49 |
|  | ILMN-PCR-free-3 | HG008-N-D | 0.6 | 150 | 444 | 102 | 97.1 | 99.6 | 115 |  |  |  |
|  |  | HG008-T | 0.5 | 149 | 454 | 102 | 98.3 | 99.8 | 118 | 161 | 80 | 40 |
| Element Aviti short insert 2x150 | Element-short-3** | HG008-N-D | 0.4 | 147 | 327 | 116 | 98.2 | 99.7 | 77 |  |  |  |
|  |  | HG008-N-P | 0.4 | 146 | 320 | 119 | 97.9 | 99.4 | 76 |  |  |  |
|  |  | HG008-T | 0.4 | 148 | 343 | 120 | 98.4 | 99.9 | 76 | 111 | 55 | 28 |
| Element Aviti long insert 2x150 | Element-long-1 | HG008-N-D | 0.6 | 149 | 793 | 440 | 97.2 | 99.7 | 26 |  |  |  |
|  |  | HG008-N-P | 0.6 | 149 | 880 | 459 | 97.4 | 99.6 | 19 |  |  |  |
|  |  | HG008-T | 0.5 | 149 | 767 | 433 | 98.6 | 99.9 | 22 | 31 | 15 | 8 |
| PacBio Onso 2x150 | PB-Onso-1 | HG008-N-D | 0.4 | 143 | 321 | 103 | 96.6* | 99.3* | 48 |  |  |  |
|  |  | HG008-T | 0.3 | 145 | 333 | 105 | 98.1* | 99.6* | 100 | 136 | 68 | 34 |
| Ultima UG100 300bp SE | Ultima-bulk-1 | HG008-N-D | 0.8 | 302 | NA | NA | NA | 99.9 | 232 |  |  |  |
|  |  | HG008-T | 0.8 | 294 | NA | NA | NA | 99.9 | 212 | 306 | 153 | 77 |
| Ultima UG100 (~170bp SE) ppmSeq | Ultima-ppmSeq-1 | HG008-N-D | 0.6 | 175 | NA | NA | NA | 99.8 | 160 |  |  |  |
|  |  | HG008-T | 0.5 | 165 | NA | NA | NA | 99.8 | 110 | 157 | 79 | 39 |

### PCR-free 2×150 bp Illumina Sequencing (Dataset ID: ILMN-PCR-free-2)

#### DNA Isolation

Genomic DNA (gDNA) was isolated from approximately five million non-viable HG008-T cells (batch 0823p23) and 40 mg of non-viable HG008-N-D tissue using the Revvity chemagic Prime 8 Instrument. HG008-T cells were resuspended and homogenized in the tissue lysis buffer, vortexed, and treated with RNase. The homogenate was then transferred to the chemagic Prime 8 instrument where DNA was extracted using the chemagic Prime DNA Blood 2k Kit H24 kit (PN: CMG-1497). For the normal tissue sample, the tissue was minced with a razor blade, incubated overnight in the lysis buffer with proteinase K. The following day the homogenate was subjected to an RNase treatment and transferred to the chemagic Prime 8 instrument where DNA was extracted using the chemagic Prime DNA blood extraction kit (CMG-1497). Genomic DNA mass yields for tumor and normal samples were approximately 68 and 72 ug, respectively. Resulting DNA was quantified using the PicoGreen Assay (Quant-iT™ PicoGreen™, catalog# P11496) and the quality was assessed using 1 % agarose gel (Invitrogen™ E-Gel™ Agarose Gels, catalog# G720801).

#### Library Preparation

Library preparation was performed for each sample using an in-house PCR-free protocol, with the Roche Kapa HyperPrep PCR-free Kit (PN: KK8505). Two libraries were prepared for the tumor sample from 1 ug of gDNA for each library and one library was prepared for the normal duodenal sample from 1 ug of gDNA. For both tumor and normal samples, DNA was sheared into fragments of approximately 450 to 600 bp using a Covaris E220 system.

Fragmentation was followed by purification of the fragmented DNA using double-sided bead clean up with different ratios of AMPure XP beads to remove very small and large fragments. DNA end-repair and 3’-adenylation were performed in the same reaction followed by ligation of the barcoded adapters using a set of 96 8-bp barcoded adapters from IDT, Illumina TruSeq RNA unique dual (UD) Indexes (PN: 20040870). Post ligation products were cleaned twice with AMPure XP beads to get rid of leftover adapters/adapter dimers.

#### Whole Genome Sequencing

The resulting tumor and normal PCR-free libraries were sequenced on an Illumina NovaSeq 6000 (ICS v1.8.1 and bcl2fastq2 version 2.20.0.422). The tumor libraries were pooled with other samples and loaded in a single lane across four flowcells. The normal library was sequenced in a pool of samples and loaded in one lane. Paired-end, 2×150 bp sequencing, was performed using the Illumina NovaSeq 6000 S4 Reagent Kit v1.5 - 300 cycles (PN: 20028312) with the NovaSeq XP 4-Lane Kit v1.5 (PN: 20043131).

#### Validation Methods

The Illumina sequencing paired-end FASTQs were assessed with FastQC. The Illumina paired-end FASTQ data were aligned to the GRCh38 reference genome using BWA-MEM. The GA4GH QC pipeline was run on the resulting BAM files to assess the quality metrics of the alignments (Table S2, see Supplementary Information document). A summary of QC metrics can be found Table 3.

### PCR-free 2×150 bp Illumina Sequencing (Dataset ID: ILMN-PCR-free-3)

#### DNA Isolation

Genomic DNA (gDNA) was isolated from approximately two million non-viable HG008-T cells (batch 0823p23) using Qiagen’s QIAamp Blood Mini Kit (PN: 51104), yielding approximately 6.4 ug of gDNA as measured by a Qubit Fluorometer and sizing with peak at approximately 21 kb as measured by an Agilent Fragment Analyzer. HG008-N-D gDNA was previously isolated from non-viable tissue and quality assessed upon receipt, approximately 2.5 ug as measured by a Qubit Fluorometer and sizing with peak at approximately 53 kb as measured by an Agilent Tapestation.

#### Library Preparation

PCR-free library preparation was performed using the Illumina TruSeq DNA PCR-Free High Throughput Library Prep Kit (96 samples) (PN: 20015963), starting with one microgram of gDNA sample. Genomic DNA samples were concentration normalized on a Hamilton robot. Intact gDNA was sheared using the Covaris LE220 sonicator. Fragmentation was followed by an initial cleanup and end-repair, which was then followed by a double-sided bead-based size selection of the repaired DNA fragments. Size selection was followed by A-tailing, in turn followed by ligation using unique dual indexed (UDI) adapters (HG008-T: TTCCTGTT-AAGATACT, HG008-N-D: CGGACAAC-AATCCGGA) to prevent index hopping on patterned flow cells. Two final bead-based clean-ups of the libraries were subsequently performed. The uniquely indexed libraries were then validated using the Fragment Analyzer (Agilent) and pooled based on concentration as determined by Picogreen (ThermoFisher). The final libraries were evaluated for size distribution on the Fragment Analyzer and are quantified by qPCR with adapter specific primers (Kapa Biosystems).

#### Whole Genome Sequencing

The tumor and normal TruSeq libraries were prepared and sequenced approximately four months apart. A single library was prepared for each sample. For each tumor and normal, the TruSeq library was multiplexed and loaded across four lanes of three flow cells to achieve the desired coverage. Paired-end, 2×150 bp sequencing, was performed using the Illumina S4 Reagent Kit v1.5 (300 cycles) (PN: 20028312) on an Illumina NovaSeq 6000 (ICS v1.7.5, RTA v3.4.4).

#### Validation Methods

The Illumina sequencing paired-end FASTQs were aligned to the GRCh38 reference genome using BWA-MEM as part of the NYGC somatic cancer pipeline (https://bitbucket.nygenome.org/projects/WDL/repos/somatic_dna_wdl/browse/README_pipeline.md?at=refs%2Fheads%2Fgiab#deliverables). Prior to alignment, pre-processing of the data was performed including adapter trimming and alignment included short-alignment marking, Novosort duplicate marking, GATK Base quality score calibration (BSQR). The GA4GH QC pipeline was run on the resulting BAM files to assess the quality metrics of the alignments (Table S2, see Supplementary Information document). A summary of QC metrics can be found in Table 3.

### Element Aviti

#### Short and long insert PCR-free 2×150 bp Aviti Sequencing (Dataset IDs: Element-short-3, Element-long-1)

##### Standard (short) Insert Library Preparation

Library preparation was performed using previously isolated genomic DNA (gDNA), as described above in “Tumor and normal samples for measurement”, from non-viable HG008-T cells (batch 0823p23) and non-viable HG008-N-D and HG008-N-P tissue. One short insert library (350 to 400 bp) was prepared from each gDNA source. For each library, 500 ng of gDNA was input into the Element Elevate Mechanical Library Prep Kit (PN: 830-00008, Document # MA-00004 Rev. E) using the Elevate Long UDI Adapter Kit Set A (PN: 830-00010). Mechanical shearing of gDNA was performed using a Covaris ultrasonicator to achieve the desired insert size of approximately 400 bp. For the short insert libraries seven iterations with the following settings were performed: 10 second duration, peak power 50, duty cycle 20, 1000 cycles per burst, average power 10. Shearing was followed by end repair and A-tailing. PCR-free library preparation continued with adapter ligation, generating dual-index libraries. Size selection was performed with double-sided SPRI bead clean-up with SPRI ratios of 0.46X/0.62X. Linear short insert libraries were quantified for loading onto the AVITI system using qPCR.

##### Long Insert Library Preparation

Library preparation was performed using previously isolated genomic DNA (gDNA) from non-viable HG008-T cells (batch 0823p23), as described above in “Tumor and normal samples for measurement”, and non-viable HG008-N-D and HG008-N-P tissue. One long insert library (approximately 900 bp) was prepared from each gDNA source. For each library, one microgram of gDNA was input into the Element Elevate Mechanical Library Prep Kit (PN: 830-00008, Document # MA-00004 Rev. E) using the Elevate Long UDI Adapter Kit Set A (PN: 830-00010). Mechanical shearing of gDNA was performed using a Covaris ultrasonicator to achieve a longer insert size of approximately 800 to 900 bp. For the long insert libraries one iteration with the following settings were performed: 5 second duration, peak power 50, duty cycle 20, 1000 cycles per burst, average power 10. PCR-free library preparation continued with end-repair, A-tailing and adapter ligation, generating dual-index libraries. Size selection was performed with double-sided SPRI bead clean-up with SPRI ratios of 0.3X/0.42X. Linear libraries were quantified for loading onto the AVITI system using qPCR using the 2x Phanta Flash master mix (Vazyme PN: P510) and Evagreen (Biotium PN:31000).

##### Whole Genome Sequencing

Linear libraries were loaded onto individual high output flowcells, 6 flowcells in total (3 for short insert, 3 for long insert). The Element AVITI 2×150 Sequencing Kit Cloudbreak UltraQ sequencing chemistry (PN: 860-00018) was used to generate 150 bp paired-end reads on the AVITI system.^10^

##### Validation Methods

The Element AVITI UltraQ paired-end FASTQs were assessed using FastQC. The FASTQs were aligned to the GRCh38 reference genome using BWA-MEM. The GA4GH QC pipeline was run on the alignment BAM files to assess the quality metrics of the alignments (Table S2, see Supplementary Information document). A summary of QC metrics can be found Table 3.

### PacBio Onso

#### PCR-free 2×150 bp Onso Sequencing (Dataset ID: PB-Onso-1)

##### Library Preparation

Library preparation was performed using previously isolated genomic DNA (gDNA), as described above in “Tumor and normal samples for measurement”, from non-viable HG008-T cells (batch 0823p23) and non-viable HG008-N-D tissue. PCR-free library preparation was performed using the PacBio Onso fragmentation DNA library prep kit (PN: 102-499-100). Two libraries were prepared, one for each of the tumor (HG008-T) and normal duodenal (HG008-N-D) samples starting with 200 ng of high molecular weight (HMW) gDNA. HG008-T and HG008-N-D HMW gDNA were first enzymatically fragmented followed by end-repair, A-tailing and adapter ligation per the fragmentation kit protocol. A small aliquot of libraries were amplified using the PacBio Onso library amp kit (PN: 102-410-800) for library QC purposes (library size measurement by Agilent TapeStation). Quantitative PCR was performed with the PacBio Onso library quant kit (PN: 102-431-800) to quantify Onso-compatible libraries for flow cell loading.

##### Whole Genome Sequencing

Whole genome sequencing of the resulting PCR-free libraries was performed on a PacBio Onso system using sequencing-by-binding (SBB) technology.^11^ The tumor library was loaded across four lanes of two flow cells and the normal duodenal library across two lanes of one flow cell. Cluster generation and paired-end 2×150 bp reads were generated using the PacBio Onso 300 cycle sequencing kit (PN: 102-860-300). Onso reverse (R2) reads may contain 1 bp reads. It is advised these reads be filtered out prior to further processing using a suitable trimmer such as cutadapt.^12^

##### Validation Methods

The PacBio Onso paired-end FASTQs were assessed with FastQC. The FASTQs were aligned to the GRCh38 reference genome using BWA-MEM. The GA4GH QC pipeline was run on the resulting BAM files to assess the quality metrics of the alignments (Table S2, see Supplementary Information document). A summary of QC metrics can be found Table 3.

### Ultima UG 100

#### PCR-free 300 bp single-end UG 100 Sequencing (Dataset ID: Ultima- bulk-1)

##### Library Preparation

Library preparation was performed using previously isolated genomic DNA (gDNA), as described above in “Tumor and normal samples for measurement”, from non-viable HG008-T cells (batch 0823p23) and non-viable HG008-N-D tissue. Library preparation for each sample, started with 500 ng of isolated genomic DNA. PCR-free library preparation was performed using the NEBNext® Ultra™ II FS DNA PCR-free Library Prep Kit (PNs: E7805, E6177) following Ultima Genomics NEB PCR-free WGS Library Preparation Protocol (Publication #: P00014 Rev. D). Ultima Genomics PCR-free adapters, xGen™ PCR-free Adapters for Ultima P1, 96 rxn (PN: 10016841) for UG 100 Sequencer compatibility were used.

##### Whole Genome Sequencing

Sequencing was performed on the UG 100 Sequencer (software version: APL 5.1.0.19) using iV27 chemistry.^13^ Single-end, 300 bp reads were generated for both samples using the UG_116cycles_Baseline_1.5.3.2 sequencing recipe and flow order [T,G,C,A] on a single wafer. A human genome control sample HG002 (Coriell Nucleic acid ID: NA24385) was spiked into each run in order to ensure quality of the samples being sequenced.

##### Validation Methods

The Ultima single-end FASTQs were aligned to the GRCh38 reference genome using the optimized version of the BWA developed by Ultima Genomics. The CRAM files were evaluated using the GA4GH Quality control (QC) pipeline to assess the quality metrics of the alignments. Alignment and validation tools can be found in Table S2, see Supplementary Information document. A summary of QC metrics can be found Table 3.

### ppmSeq UG 100 Sequencing (Dataset ID: Ultima-ppmSeq-1)

#### Library Preparation

Library preparation was performed using previously isolated genomic DNA (gDNA), as described above in “Tumor and normal samples for measurement”, from non-viable HG008-T cells (batch 0823p23) and non-viable HG008-N-D tissue. For each sample, one paired plus minus sequencing (ppmSeq) library was prepared starting with either 180 ng of HG008-T gDNA or 250 ng of isolated HG008-N-D gDNA. PpmSeq library preparation was performed using the NEBNext® Ultra™ II DNA PCR-free Library Prep Kit (PN: E7410) following the Ultima ppmSeq protocol (UG ppmSeq Genomic DNA Library preparation Protocol for Solaris Free D100093 Rev.02, see Supplementary File 1), UG ppmSeq Adapters (96-sample indices plate, PN: UG0105) for UG 100 Sequencer compatibility were used.

#### Whole Genome Sequencing

Sequencing was performed on the UG 100 Sequencer (software version: APL 5.1.0.19) using iV27 chemistry. A human genome control sample HG002 (Coriell Nucleic acid ID: NA24385) was spiked into each run in order to ensure quality of the samples being sequenced. Libraries were pooled and single-end, average 165 bp (tumor) and 175 bp (normal) reads were generated for both samples using the UG_116cycles_Baseline_1.7.0.0 sequencing recipe and flow order [T,G,C,A] on a single wafer.

#### Validation Methods

Ultima ppmSeq reads (single end) which are output of dsDNA beads contain the information of both strands (i.e. duplex). These reads are referred to as “Mixed” reads. Typically, around 40%-50% of the sequenced reads are Mixed. In this experiment, the Mixed reads coverage was 53.12X on average for the tumor sample (48%) and 76.14X for the normal sample (44%). The Ultima single-end FASTQs were aligned to the GRCh38 reference genome using the optimized version of the BWA developed by Ultima Genomics. The CRAM files were evaluated using the GA4GH Quality control (QC) pipeline to assess the quality metrics of the alignments. Alignment and validation tools can be found in Table S2, see Supplementary Information document. A summary of QC metrics can be found Table 3.

### Hi-C library Preparation

#### Phase Genomics with Illumina sequencing (Dataset ID: HiC-ILMN-1 and HiC-ILMN-3)

##### Library Preparation

CytoTerra library preparation was performed using approximately 500,000 viable HG008-T cells from passage 38 of the MGH 2022 culture and 250,000 viable HG008-T cells from passage 21 of the NIST 2024 culture for HiC-ILMN-1 and HiC-ILMN-3, respectively. Proximity ligation preparation was performed with the Phase Genomics Proximo Hi-C v4.5 kit (PN: KT4045) and associated Proximo human protocol (v4.5 0202301, https://4408603.fs1.hubspotusercontent-na1.net/hubfs/4408603/Protocols_latest/Proximo_Human_v4.5_202301.pdf) to generate a dual-indexed Illumina-compatible libraries.^14^ In brief, intact cells are crosslinked, lysed, and chromatin digested with a cocktail of restriction endonucleases. Ends are filled in with biotinylated nucleotides and subjected to proximity ligation. Following ligation, crosslinks are enzymatically reversed and DNA purified. Proximity ligated junctions are purified using streptavidin beads. Bead-bound DNA is used as input for generation of a paired-end Illumina compatible library.

##### Whole Genome Sequencing

The Proximo Hi-C libraries were run on either an Illumina NovaSeq 6000 (NCS v1.7.5, RTA v3.4.4) for HiC-ILMN-1 libraries or a NovaSeq X (NCS 1.2.2.48004) for the HiC-ILMN-3 libraries. The libraries were sequenced using a 2×150bp paired-end configuration using associated Illumina instrument compatible kits (NovaSeq 6000 S4 Reagent Kit v1.5 300 cycles PN: 20028312, NovaSeq X Series 10B Reagent Kit PN: 20085594). For each run the associated libraries were loaded on a single lane of a flowcell. Image analysis and base calling were conducted by the NovaSeq Control Software. Raw sequence data (.bcl files) generated from the Illumina NovaSeq was converted into FASTQ files and de-multiplexed using Illumina bcl2fastq 2.20 software. One mis-match was allowed for index sequence identification.

##### Validation Methods

QC of the HiC data was performed as part of the documented Phase Genomics alignment and QC process (https://phasegenomics.github.io/2019/09/19/hic-alignment-and-qc.html). Reads were aligned to the GRCh38 reference genome using BWA-MEM and resulting alignment QC performed by running the HiC QC python script, hic_qc.py (Table S2, see Supplementary Information document). QC metrics, used for validation, are further discussed in the Technical Validation section.

### Arima with Illumina sequencing (Dataset ID: HiC-ILMN-2)

#### HiC Sample Preparation

Arima HiC library preparation used the Arima HiC High Coverage Kit (PN: A410110) following the associated User Guide for Mammalian Cell Lines (PN: Doc A160161 v01, https://arimagenomics.com/wp-content/files/User-Guide-Arima-High-Coverage-for-Mammalian-Cell-Lines.pdf).^15^ For each of the HG008-T and HG008-N-D samples, approximately one million cells were input into the HiC protocol for crosslinking chromatin. The crosslinked chromatin is then digested using a restriction enzyme cocktail optimized for coverage uniformity across a wide range of genomic sequence compositions. The 5’-overhangs are then filled in, causing the digested ends to be labeled with a biotinylated nucleotide. Next, spatially proximal digested ends of DNA are ligated, capturing the sequence and structure of the genome. The ligated DNA is then purified, producing pure proximally-ligated DNA. The proximally-ligated DNA is then fragmented, and the biotinylated fragments are enriched. HiC ligation resulted in 871 ng HiC ligation product for HG008-T and 1760 ng of HiC ligation product for HG008-N-D.

#### Library Preparation

The HG008-T and HG008-N-D HiC ligation products were prepared for Illumina Sequencing using the Roche KAPA Hyper Prep kit (PN: 7962347001) following the KAPA Hyper Prep kit User Guide (PN: A160139 v00, https://arimagenomics.com/wp-content/files/User-Guide-Library-Preparation-using-KAPA-Hyper-Prep-Kit.pdf). Library preparation begins with DNA fragmentation, DNA size selection, and biotin enrichment. For each sample, 250 ng of sheared and size-selected ligation products were input into a custom Arima-HiC end-repair, dA-tailing and adapter ligation is performed as described in the User Guide.

#### Whole Genome Sequencing

Libraries were shipped to the Baylor College of Medicine Human Genome Sequencing Center (HGSC) for sequencing. The tumor and normal samples were sequenced in a pool of samples in one lane. Paired-end, 2×150 bp sequencing, was performed using the Illumina S4 Reagent Kit v1.5 (300 cycles) (PN: 20028312) on an Illumina NovaSeq 6000 (ICS v1.7.5, RTA v3.4.4). The resulting data is referred to as Arima High Coverage HiC data.

#### HiC Validation

QC was performed as part of the Arima SV pipeline (v1.3, https://github.com/ArimaGenomics/Arima-SV-Pipeline). The pipeline, described here, https://arimagenomics.com/wp-content/files/Bioinformatics-User-Guide-Arima-Structural-Variant-Pipeline.pdf, will run HiCUP^16^ to align the Illumina sequencing reads of the Arima Hi-C library to the GRCh38 reference genome (Table S2, see Supplementary Information document). The pipeline also produces structural variant calls and visualizations however, are outside the scope of this paper. HiC data QC statistics are then generated for Illumina deeply sequenced data. QC metrics, used for validation, are further discussed in the Technical Validation section.

### Dovetail with Illumina sequencing (Dataset ID: HiC-ILMN-4)

#### Library Preparation

LinkPrep library preparation was performed starting with 1×10^6^ non-viable HG008-T cells (2024 NIST passage 21) using the Dovetail Genomics LinkPrep kit (21025V, v2.0, https://cantatabio.com/wp-content/uploads/2025/02/Dovetail_LinkPrep_Kit_User_Guide_version-2.0_Mammals-1.pdf).^17^ The cells were prepared following the mammalian cell line sample preparation protocol (Stage 1A) for crosslinking and in situ tagmentation. Proximity ligation, crosslink reversal and DNA purification was then performed. Purified crosslinked reversed DNA from the 1×10^6^ cells was split evenly for four separate library preparations. Libraries were dual-indexed, purified and size selected for the final library size range of 350 - 1,000 bps.

#### Whole Genome Sequencing

The resulting four LinkPrep libraries were then pooled for sequencing on an Illumina NovaseqXplus with 2×150 paired end reads using the Illumina NovaSeq™ X Series 25B Reagent Kit (300 Cycle) (PN: 20104706). The pooled library was loaded onto one lane of a single flowcell.

#### Validation Methods

The HG008-T LinkPrep library QC followed the Alignment and Proximity-Ligation QC process (https://varilink.readthedocs.io/en/latest/library_qc.html) to report pairing statistics. The QC process aligned the reads to the GRCh38 reference genome using BWA-MEM (v2.2.1) and retained high quality mapped reads. The mapped data was then used to generate a pairs file with pairtools, which categorizes pairs by read type and insert distance, this step both flags and removes PCR duplicates. Once pairs are categorized, counts of each class are summed and reported. QC metrics, used for validation, are further discussed in the Technical Validation section.

### Bulk long-read whole genome sequencing**^8^**

#### PacBio Revio HiFi

##### PCR-free Revio HiFi Sequencing (Dataset ID: PB-HiFi-1)

###### Library Preparation

Library preparation was performed using previously isolated genomic DNA (gDNA), as described above in “Tumor and normal samples for measurement”, from non-viable HG008-T cells (batch 0823p23) and non-viable HG008-N-P tissue. Sizing of the isolated gDNA was performed prior to library preparation to confirm the gDNA was of sufficient length for long-read sequencing. The average sizing of smear analysis from Pulsed Field Gel Electrophoresis (PFGE) with an Femto Pulse System (Agilent) was approximately 58 kbp for HG008-T and 29 kbp for HG008-N-P.

Library preparation was performed following PacBio procedure and checklist: “Preparing whole genome and metagenome libraries using SMRTbell® prep kit 3.0” (PN: 102-166-600 REV02 MAR2023). This protocol uses the PacBio SMRTbell prep kit 3.0 (PN: 102-182-700), SMRTbell adapter index plate 96A (PN: 102-009-200), and SMRTbell cleanup bead kit (PN 103-306-300). Briefly, 6.4 ug of HG008-T and 6.1 ug of HG008-N-P were sheared separately on a Megaruptor 3 (Diagenode, B06010003) to a target size of 15 to 20 kb. Two library preparations were performed, one using sheared gDNA inputs of 3.1 ug of HG008-T and one using 4.1 ug of HG008-N-P. Sheared samples were subjected to a combined DNA repair and A-tailing reaction followed by ligation of sequencing adapters to create SMRTbell libraries. SMRTbell libraries were treated with a nuclease mix to remove unligated library components and then size-selected on a Sage Science PippinHT system using a 0.75 % agarose gel cassette with cassette definition file, “6–10kb High Pass Marker 75E” to remove fragments less than 10 kb.

###### Whole Genome Sequencing

PacBio SMRTbell libraries were prepared for HiFi sequencing (i.e. circular consensus sequencing)^18^ by annealing and binding sequencing primers and polymerase, respectively, to the SMRTbell templates using Revio polymerase kit (PN 102-817-600) and following instructions provided in the Sample Setup module of PacBio SMRT Link software (v13.0), a web-based end-to-end sequencing workflow manager. Sequencing-ready SMRTbell libraries were sequenced on a Revio system using the following reagents and consumables: Revio sequencing plate (PN: 102-587-400) and Revio SMRT Cell tray (PN: 102-202-200). 24-hour movies were run. Two SMRT Cells were run for HG008-T and one SMRT Cell for HG008-N-P.

###### Validation Methods

The PacBio Revio unaligned bams were aligned to the GRCh38 reference genome using the PacBio pbmm2 aligner as part of the PacBio HiFi-human-WGS_WDL (germline) pipeline v1.1.0 (https://github.com/PacificBiosciences/HiFi-human-WGS-WDL). Cramino, Nanoplot and Samtools stats tools were used to assess the quality of the alignments (Table S2, see Supplementary Information document). A summary of QC metrics can be found Table 4. While the method used maintains base modification (methylation) information, this was not assessed as part of the data validation process.

**Table 4.** Long-read metrics. Summary of metrics generated from Cramino and Mosdepth for validation of PacBio HiFi and ONT data. All mapping-based metrics presented are based on alignments to the GRCh38-GIABv3 reference genome.

| Technology | Dataset ID | GIAB sample ID | number of alignments | percent of bases mapped | yield (Gb) | mean coverage | N50 | median identity to GRCh38 | diploid mean coverage | haploid mean coverage | NRPCC |
| --- | --- | --- | --- | --- | --- | --- | --- | --- | --- | --- | --- |
| PacBio HiFi | PB-Hifi-1 | HG008-N-P | 6,903,194 | 99.8 | 110 | 35 | 16,764 | 99.8 |  |  |  |
|  |  | HG008-T | 14,502,556 | 99.8 | 245 | 79 | 17,833 | 99.8 | 116 | 58 | 29 |
|  | PB-Hifi-2 | HG008-N-D | 13,268,351 | 99.7 | 212 | 68 | 16,658 | 99.8 |  |  |  |
|  |  | HG008-T | 12,974,130 | 99.6 | 224 | 72 | 18,132 | 99.8 | 106 | 53 | 27 |
| ONT standard | ONT-std-1 | HG008-N-D | 38,959,690 | 98.3 | 292 | 94 | 12,516 | 99.2 |  |  |  |
|  |  | HG008-N-P | 9,591,713 | 98.2 | 126 | 41 | 24,460 | 99.1 |  |  |  |
|  | ONT-std-2 | HG008-T | 6,673,103 | 99.7 | 138 | 45 | 35,497 | 99.2 | 63 | 31 | 16 |
| ONT Ultra Long | ONT-UL-1 | HG008-T | 2,576,733 | 99.6 | 110 | 39 | 118,692 | 98.7 | 54 | 27 | 14 |
|  | ONT-UL-2 | HG008-T | 4,068,842 | 99.0 | 119 | 38 | 83,765 | 99.6 | 54 | 27 | 14 |

### PCR-free Revio HiFi Sequencing (Dataset ID: PB-HiFi-2)

#### DNA Isolation

Genomic DNA (gDNA) was isolated from approximately five million non-viable HG008-T cells (batch 0823p23) and 40 mg of non-viable HG008-N-D tissue using the Revvity chemagic Prime 8 Instrument. Cells were resuspended and homogenized in the tissue lysis buffer, vortexed, and treated with RNase. The homogenate was then transferred to the chemagic Prime 8 instrument where DNA was extracted using the chemagic Prime DNA Blood 2k Kit H24 kit (PN: CMG-1497). For the normal tissue sample, the tissue was minced with a razor blade, incubated overnight in the lysis buffer with proteinase K. The following day the homogenate was subjected to an RNase treatment and transferred to the chemagic Prime 8 instrument where DNA was extracted using the chemagic Prime DNA blood extraction kit (CMG-1497). Genomic DNA mass yields for tumor and normal samples were approximately 68 and 72 ug, respectively. Resulting DNA size was measured on an Agilent Femto Pulse System with average sizing of approximately 81 kb for gDNA from tumor cell line and 59 kb for gDNA from normal duodenal tissue.

#### Library Preparation

Genomic DNA (7.5 ug) was sheared using Covaris g-tubes to achieve an average size of 15 to 20 kb. Sheared DNA was size selected with a Sage Science PippinHT using the 15 to 20 kb high-pass 75E cassette definition (gel cassette HPE7510) to eliminate fragments smaller than 15 kb. One library for each sample type was prepared. The PacBio procedure for library preparation was performed using the PacBio SMRTbell prep kit 3.0 (PN: 102-182-700) and the SMRTbell adapter index plate 96A (PN: 102-009-200) for repair and A-tailing, adapter ligation and nuclease treatment to remove unligated fragments. Final libraries were purified using 1X PacBio SMRTBell Cleanup beads, eluted in 30 µL PacBio elution buffer and quantified using the Qubit dsDNA quantification high-sensitivity assay (Thermo Fisher Scientific). Final library size was determined using the Agilent Femtopulse.

#### Sequencing Methods

PacBio SMRTbell libraries were prepared for HiFi sequencing (i.e. circular consensus sequencing) by annealing and binding sequencing primers and polymerase, respectively, to the SMRTbell templates using Revio polymerase kit (PN: 102-817-600) and following instructions provided in the Sample Setup module of PacBio SMRT Link software (13.0.0.207600). Sequencing-ready SMRTbell libraries were sequenced on a PacBio Revio system (ICS version 13.0.0.212033). 30-hour movies were run. Two SMRT cells each were run for HG008-T and HG008-N-D. QC metrics from SMRTlink software including HiFi reads, mean read length, P1 %, etc were as expected.

#### Validation Methods

The PacBio Revio sequencing unaligned bams raw data were aligned to the GRCh38 reference genome using the PacBio pbmm2 aligner as part of the PacBio Somatic variant calling Pipeline v0.7 (https://github.com/PacificBiosciences/HiFi-Somatic-WDL). Cramino, Nanoplot and Samtools stats tools were used to assess the quality of the alignments (Table S2, see Supplementary Information document). A summary of QC metrics can be found Table 4. While the method used maintains base modification (methylation) information, this was not assessed as part of the data validation process.

### Oxford Nanopore Technologies PromethION

#### PCR-free Standard Length PromethION Sequencing (Dataset ID: ONT-std-1)

##### Library Preparation

Library preparation was performed using previously isolated genomic DNA (gDNA), as described above in “Tumor and normal samples for measurement”, from non-viable HG008-N-P and HG008-N-D tissues. Library preparation was performed with Oxford Nanopore Technologies Ligation Sequencing Kit V14 (PN: SQK-LSK114) using one microgram of gDNA from each tissue.

##### Whole Genome Sequencing

Nanopore based sequencing was performed on Oxford Nanopore Technologies (ONT) PromethION utilizing ONTs PromethION R10.4.1 Flow Cells (PN: FLO-PRO114M).^19^ To maximize throughput multiple flow cells were used. Sequencing was performed using four flow cells for the duodenal sample and three flow cells for the pancreatic sample.

##### Basecalling

The ONT Dorado v0.5.3 basecaller (https://github.com/nanoporetech/dorado) and DNA model, ‘dna_r10.4.1_e8.2_400bps_sup@v4.3.0_5mC_5hmC@v1’ were used for basecalling, including modified bases. Basecalling successfully completed with no errors. Output from the basecaller is a bam file of unaligned reads including modified bases.

##### Validation Methods

The Dorado basecaller generates a ‘summary.txt’ file, which was analyzed using PycoQC to assess sequencing quality metrics. The ONT unaligned bams were processed to extract the FASTQ sequences by using samtools then aligned to the GRCh38 reference genome using the Minimap2. Cramino, Nanoplot and Samtools stats tools were used to assess the quality of the alignments (Table S2, see Supplementary Information document). A summary of QC metrics can be found Table 4. While the method used maintains base modification information, this was not assessed as part of the data validation process.

### PCR-free Standard Length PromethION Sequencing (Dataset ID: ONT-std-2)

#### DNA Isolation

DNA was extracted from approximately five million non-viable HG008-T cells (batch 0823p23) utilizing the NEB Monarch® HMW DNA Extraction Kit for Tissue (PN: T3060). Total DNA extracted was approximately 80 µg. Only 5 µg of DNA was sheared using Diagenode’s Megaruptor3 (PN: B06010003) with the DNAFluid+ Kit (PN: E07020001) followed by DNA size selection using PacBio SRE kit (PN: 102-208-300). The size of sheared DNA fragments were analyzed on the Agilent Femto Pulse System using genomic DNA 165 kb kit (PN: FP-1002-0275). Fragment size distribution of post-sheared DNA had a peak at approximately 45 kb length.

#### Library Preparation

The UCSC protocol has been optimized to generate N50s of approximately 30 kb with very high throughput for ligation-based sequencing. Library preparation was performed with Oxford Nanopore Technologies (ONT) Ligation Sequencing Kit XL V14 (PN: SQK-LSK114-XL). Enough library was created for 4 library loads onto flow cells.

#### Whole Genome Sequencing

Nanopore based sequencing was performed on the ONT PromethION 48/PromethION A-Series Data Acquisition Unit (PN: PRO-SEQ048/PRO-PRCAMP) utilizing ONT PromethION R10.4.1 Flow Cells (PN: FLO-PRO114M). Sequencing was performed on the same day as ligation library preparation. Sequencing was performed utilizing one PromethION R10.4.1 Flow Cell. The flow cell was washed with ONTs Flow Cell Wash Kit XL (PN: EXP-WSH004-XL) each day and re-primed and loaded with fresh library for 4 days of sequencing to maximize throughput.

#### Basecalling

The ONT Dorado v0.3.4 basecaller (https://github.com/nanoporetech/dorado) and model ‘dna_r10.4.1_e8.2_400bps_sup@v4.2.0/dna_r10.4.1_e8.2_400bps_sup@v4.2. 0_5mCG_5hmCG@v2’ were used for basecalling, including modified bases. Basecalling successfully completed with no errors. Output from the basecaller is a bam file of unaligned reads including modified bases.

#### Validation Methods

The Dorado basecaller generates a ‘summary.txt’ file, which was analyzed using PycoQC to assess sequencing quality metrics. The Oxford Nanopore Technologies sequencing unaligned bams were processed to extract the FASTQ sequences then aligned to the GRCh38 reference genome using the Minimap 2. Cramino, Nanoplot and Samtools stats tools were used to assess the quality of the alignments (Table S2, see Supplementary Information document). A summary of QC metrics can be found Table 4. While the method used maintains base modification (methylation) information, this was not assessed as part of the data validation process.

### PCR-free Ultra Long PromethION Sequencing (Dataset ID: ONT-UL-1)

#### DNA Isolation

Ultra-high molecular weight DNA was extracted from approximately five million non-viable HG008-T cells (batch 0823p23) utilizing New England Biolabs Monarch® HMW DNA Extraction Kit for Tissues (NEB #T3060). Wide bore pipette tips were used throughout to preserve ultralong DNA. No QC of resulting gDNA was performed as viscosity of isolated DNA solution is challenging to work with. Rather ultra long DNA is assessed as part of the Oxford Nanopore Technologies (ONT) sequencing and previous experience with a five million cell as input.

#### Library Preparation

Library preparation was performed utilizing the ONT Ultra-Long DNA Sequencing Kit Version 14 (SQK-ULK114). The library was eluted in approximately 810 µL volume in order to sequence across additional flow cells to maximize data collection.

#### Whole Genome Sequencing

Nanopore based sequencing was performed on an ONT PromethION 48/PromethION A-Series Data Acquisition Unit (PN: PRO-SEQ048/PRO-PRCAMP) utilizing ONT PromethION R10.4.1 Flow Cells (PN: FLO-PRO114M). Sequencing was performed the day following library prep utilizing three PromethION R10.4.1 Flow Cells. The flow cells were washed with ONTs Flow Cell Wash Kit XL (EXP-WSH004-XL) each day and reloaded with the library for 4 days of sequencing to maximize throughput.

#### Basecalling

The ONT Dorado v0.4.3 basecaller (https://github.com/nanoporetech/dorado) and model, ‘dna_r10.4.1_e8.2_400bps_sup@v4.2.0/dna_r10.4.1_e8.2_400bps_sup@v4.2. 0_5mCG_5hmCG@v2’ were used for basecalling, including modified bases. Basecalling successfully completed with no errors. Output from the basecaller is a bam file of unaligned reads including modified bases.

#### Validation Methods

The Dorado basecaller generates a ‘summary.txt’ file, which was analyzed using PycoQC to assess sequencing quality metrics.The ONT unaligned bams were processed to extract the FASTQ sequences by using samtools then aligned to the GRCh38 reference genome using the Minimap2. Cramino, Nanoplot and Samtools stats tools were used to assess the quality of the alignments (Table S2, see Supplementary Information document). A summary of QC metrics can be found Table 4. While the method used maintains base modification (methylation) information, this was not assessed as part of the data validation process.

### PCR-free Ultra Long PromethION Sequencing (Dataset ID: ONT-UL-2)

#### DNA Isolation

Ultra-high molecular weight DNA was extracted from approximately five million non-viable HG008-T cells (batch 0823p23) utilizing New England Biolabs Monarch® HMW DNA Extraction Kit for Cells (NEB #T3050).

Extraction yielded ∼25-30 ug with DNA hundreds of kilobases long, with some exceeding megabases, as measured by preliminary analysis of Oxford Nanopore Technologies (ONT) sequencing.

#### Library Preparation

A single library preparation was performed utilizing the ONT Ultra-Long DNA Sequencing Kit Version 14 (SQK-ULK114) using all isolated gDNA.

#### Whole Genome Sequencing

Nanopore based sequencing was performed on an ONT 48/PromethION A-Series Data Acquisition Unit (PN: PRO-SEQ048/PRO-PRCAMP) utilizing ONTs ULK114 chemistry with the E8.2.1 motor protein and PromethION R10.4.1 Flow Cells (PN: FLO-PRO114M). Two flow cells were used and received a total of four library loads with three wash (ONT Flow Cell Wash Kit XL PN: EXP-WSH004-XL) and reload steps per flowcell.

#### Basecalling

The ONT Dorado v0.8.1 basecaller (https://github.com/nanoporetech/dorado) and the super accuracy (sup) model (v5.0.0), were used for basecalling, including 5mC_5hmC modification calls (v2.0.1). Basecalling successfully completed with no errors. Output from the basecaller is a bam file of unaligned reads including modified bases.

#### Validation Methods

The ONT unaligned bams were processed to extract the FASTQ sequences by using samtools then aligned to the GRCh38 reference genome using the Minimap2. Cramino, Nanoplot and Samtools stats tools were used to assess the quality of the alignments (Table S2, see Supplementary Information document). A summary of QC metrics can be found Table 4. While the method used maintains base modification information, this was not assessed as part of the data validation process.

### Single cell whole genome sequencing**^8^**

#### BioSkryb ResolveDNA with Illumina Low Pass Sequencing (Dataset ID: sc-ILMN-1)

##### Whole Genome Amplification (WGA)

Approximately two million non-viable HG008-T cells (batch 0823p23) were sorted with a Sony SH800 using a 130 micron chip. Singlet (FSC-A / FSC-H, BSC-A / BSC-W) and live-cell (PI negative, top 70 % Calcein-AM positive) gating was employed for single cell sorting into a 384-well plate with 3 µL of BioSkryb Cell Buffer (PN: 100199). Individual cells were processed within wells using the ResolveDNA WGA Kit (PN: 100136) following steps outlined in the ResolveDNA Whole Genome Amplification Kit v2.0 User Guide. Primary Template Directed Amplification (PTA) to amplify single cell whole genomes and transform into Illumina-sequenceable libraries.^20^ Approximately 70 % of cells met specifications for BioSkryb WGA metrics. Dual-indexed Illumina-compatible libraries were generated for the top performing 120 cells.

##### Illumina Low-Pass Sequencing

Illumina low pass sequencing performed to QC the single cell libraries. libraries were pooled and sequenced on a single flowcell targeting 2 million reads as an initial QC using an Illumina NextSeq 1000 instrument with 2×50 paired-end reads with P2 reagents v3 (PN: 200446811).

##### Single Cell Library Validation

Single-cell libraries were assessed by running BioSkryb’s BJ-DNA-QC pipeline (https://docs.basejumper.bioskryb.com/pipelines/secondary/bj-dna-qc/docs/) through the BioSkryb analytics platform, BaseJumper. The BJ-DNA-QC pipeline uses the Illumina low-pass sequencing data and generates several QC metrics that help assess whether the single-cell libraries are ready for high-depth sequencing (Table S2, see Supplementary Information document). Briefly, reads were aligned to the GRCh38 reference and QC-metrics obtained from the derived alignment and are used to censor single-cells. QC metrics, used for validation, are further discussed in the Technical Validation section.

### BioSkryb ResolveDNA with Ultima High Throughput Sequencing (Dataset ID: sc-Ultima-1)

#### Library Conversion

BioSkryb Genomics previously prepared 120 ResolveDNA single cell libraries for Illumina low-coverage sequencing; described above for the sc-ILMN-1 dataset. For high-coverage Ultima sequencing these libraries were then converted to a format compatible with the Ultima Genomics sequencing platform (UG 100), following the Ultima Genomics Library Conversion protocol in UG Library Amplification Kit v3 User Guide (Publication #: P00025 Rev. B). Library Amplification followed option 2 for starting with non-UG adapters. The UG Library Amplification Kit v3 (supplied by Ultima Genomics) and the IDT kit xGen™ Indexing Primers for Ultima (PNs: 10016992 and 10016993) were used for conversion and enabled multiplexing of the single cell libraries. The library amplification step was modified to replace the UG Index Primer and UG Universal primer with 5.0 µL of primer mix from the appropriate well of the IDT index primer plate. Ten nanograms of input library was used for each reaction and seven cycles of PCR were performed per the amplification protocol. Final converted libraries were QC’d for size and yield using an Agilent Tapestation and Qubit, respectively. Two plates of 120 individual Ultima single-cell libraries were sent to Ultima Genomics for sequencing.

#### Whole Genome Sequencing

Sequencing of the converted (Illumina to Ultima) libraries was performed on the UG 100 (software version: APL 5.1.0.19) using V27 Chemistry. Single end, 300 bp reads were generated using the UG_116cycles_Baseline_1.5.3.2 sequencing recipe and flow order [T,G,C,A] on a single wafer. A human genome control sample HG002 (Coriell Nucleic acid ID: NA24385) was spiked into each run in order to ensure quality of the samples being sequenced.

#### Validation

To generate sequencing measures and QC statistics, a custom Ultima tool was run that implements select picard metrics (https://github.com/broadinstitute/picard). This tool collects the metrics on board of the UG100 sequencing platform in parallel with producing sorted, duplicate-marked GRCh38 aligned CRAM files. Alignments were produced using an Ultima-optimized version of BWA-MEM (Table S2, see Supplementary Information document). QC metrics, used for validation, are further discussed in the Technical Validation section.

### Whole genome optical mapping**^8^**

#### Optical Mapping with Bionano Saphyr (Dataset ID: Bionano-1)

##### DNA Isolation

Approximately two million non-viable HG008-T cells (batch 0823p23), were washed with Bionano Stabilizing Buffer (PN: 20394 and 20397) by MGH at the time of pelleting and freezing. Upon receipt of the cell pellet, ultra-high molecular weight genomic DNA (gDNA) was isolated using the Bionano SP-G2 Blood & Cell Culture DNA Isolation Kit (PN: 80060) with a yield of approximately 6.5 ug of gDNA (100 ng/µL in 65 µL elution) with an N50 greater than 150 kbp.

##### Sample Preparation

Sequence-specific labeling of the gDNA was performed using the Bionano Direct Label and Stain-G2 (DLS-G2) Kit (PN: 80046).

##### Optical Genome Mapping

Labeled DNA was loaded onto a Bionano Saphyr Chip G3.3 and measured with the Saphyr System (ICS v5.3.23013.1). Molecules and labels imaged on the Saphyr instrument were digitized and saved in a BNX format (raw data) (https://bionanogenomics.com/wp-content/uploads/2018/04/30038-BNX-File-Format-Specification-Sheet.pdf) which served as input into downstream analyses. The Rare Variant Pipeline (RVP) (https://bionano.com/wp-content/uploads/2024/04/CG-30375-Analysis-Quick-Start-Annotated-Rare-Variant-Structural-Variant-Calling.pdf) detected SVs by the “split-read” analysis, where initial molecule alignment and molecule extension refinement allows identification and detection of somatic variants, and identifies variants that are 5 kbp or longer in size. The Access software (v1.8) was used to generate high confidence callsets for SVs and unique SVs not present in the Bionano internal control database. High confidence CNV calls were generated after recentering based on the hypodiploid tumor genome. Regions where unique SV and CNV calls overlap with Bionano custom pan cancer gene list are also reported.

##### Validation

This sample was processed based on manufacturer protocol^21^ (Table S2, see Supplementary Information document). QC metrics, used for validation, are further discussed in the Technical Validation section.

### Cytogenetic analysis

#### G-Banded Karyotyping (MGH p31 karyotyping-1 and NIST p18 karyotyping-2)^34,35^

Karyotyping was performed by KaryoLogic Inc (Durham, NC) for two passages of HG008-T as shown in Figure 2. Approximately one million viable HG008-T cells were plated in T25 cell culture flasks, two from MGH and one from NIST, and sent for karyotyping. Cytogenetic analysis was performed on twenty-five G-banded metaphase spreads of the HG008-T cell line from MGH passage 31. A second analysis was performed in 2024 using viable HG008-T cells from NIST passage 18. Cytogenetic analysis was performed on twenty G-banded metaphase spreads for the HG008-T NIST passage 18 cells (2024).

**Figure 2.**
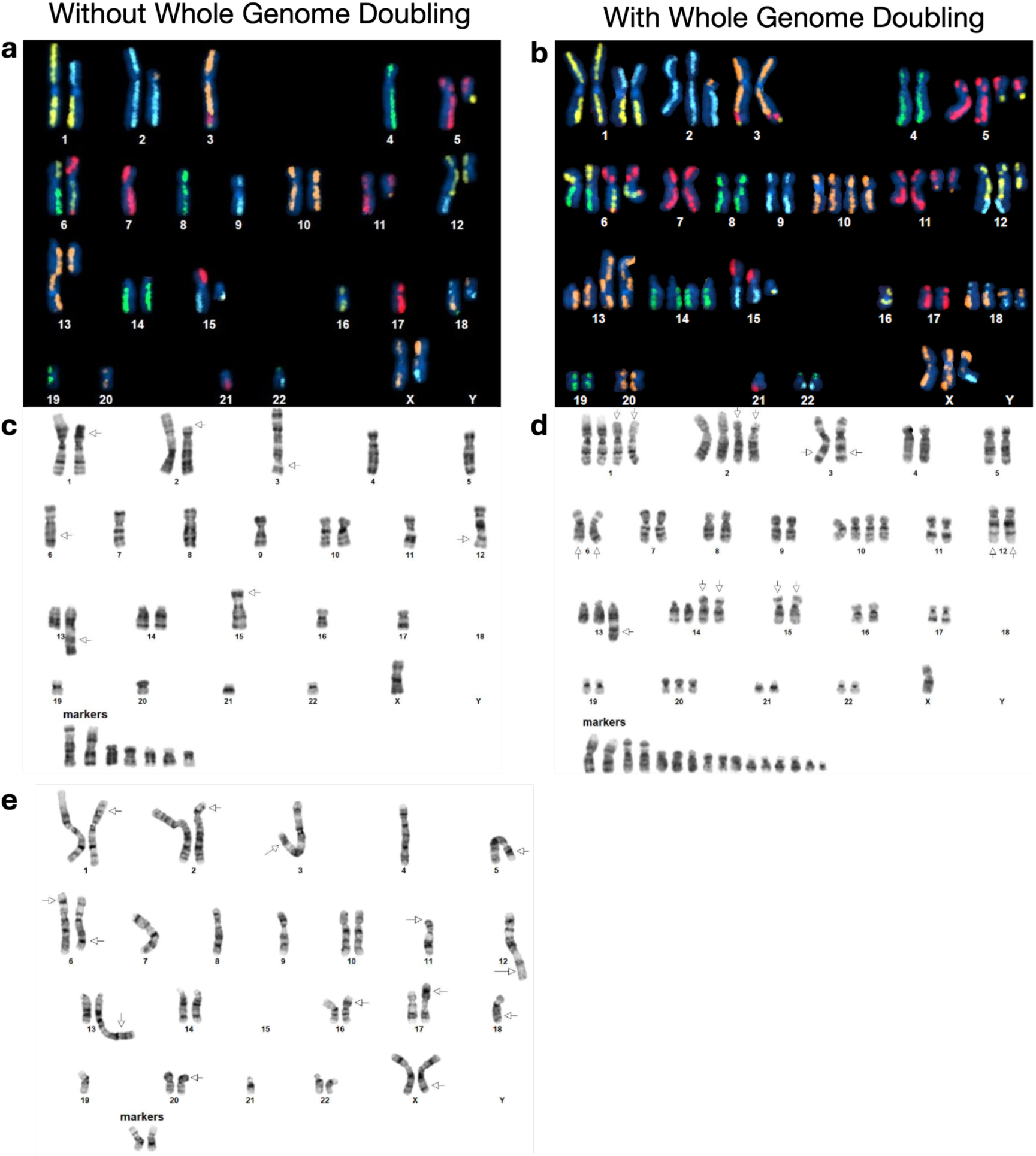
Directional Genomic Hybridization (dGH) and G-banded karyograms for three passages of the HG008-T tumor cell line. Karyograms show examples of cells with and without whole genome doubling. **(a-b)** dGH karyograms from NIST passage 21 (2024) have each chromosome colored by one of five dyes, and information from G-banded karyograms and copy number alterations were used to make preliminary assignments of chromosomes. **(c-d)** G-banded karyotypes from MGH passage 31 (2022) with and without whole genome doubling. In 25 spreads, the chromosome count varied from 29 to 71 chromosomes per spread, with the exception of two polyploid spreads of over 100 chromosomes each. While the ploidy number varied, the karyotype was relatively consistent with minor variations from spread to spread. **(e)** G-banded karyotype from NIST passage 18 (2024) without whole genome doubling, representative of 17 of 20 spreads that had 34 to 36 chromosomes. While not shown, three out of twenty spreads showed whole genome doubling, with counts ranging from 68 to 70 chromosomes.

### KROMATID Whole Genome Directional Genomic Hybridization (dGH) (Dataset ID: dGH-1)

#### Sample Preparation

Sample preparation for dGH analysis was performed at NIST from passage 21 of HG008-T. One million cells were plated in a T25 flask and were cultured overnight. The cell culture medium was replaced with dGH cell prep kit (KROMATID, Cat# dGH-0001) contained culture medium, and additional dose of dGH cell prep kit was added 4 hr after medium replacement. Cells were arrested in the first mitosis with a 4-hour Colcemid (KROMATID, Cat# COL-001) block, harvested, fixed in freshly made fixative [3:1 methanol:acetic acid (Fisher Scientific, Waltham, MA, USA)], and shipped to KROMATID. Metaphase spread preparation along with subsequent UV and exonucleolytic treatments to selectively remove the analog-incorporated daughter strand were performed at KROMATID according to published dGH protocols.^22^

#### Directional Genomic Hybridization

For whole genome analysis, KROMATID’s KROMASURE Screen assay was hybridized to the prepared metaphase spreads, then counterstained with DAPI (Vectashield, Vector Laboratories, Newark, CA, USA) and imaged on an Applied Spectral Imaging Harmony system (Applied Spectral Imaging, Carlsbad, CA, performed USA) using a 100X objective. The KROMASURE Screen assay consists of single-stranded, unidirectional, tiled oligos designed to hybridize directly to the unique sequences in each of the 24 human chromosomes. Each chromosome specific complement of oligos is end-labeled with one of five unique spectrally differentiable fluorophores, and each of these “high-density chromosome paints” are combined such that chromosomes painted in the same color can be differentiated by size, shape, and centromere position. Prior to analysis, images of KROMASURE Screen painted metaphase spreads are qualified, processed and sorted into karyograms for rapid, consistent reading of the assay.

A combination of expected KROMASURE Screen signal patterns in the (diploid) human reference genome,^23^ the HG008-T standard G-banding karyotypes, and the copy number variant was used to construct preliminary KROMASURE Screen karyograms in 10 cells, including two (2) genome-doubled cells. The prototype karyograms will be used to generate per-chromosome attribution of inter- and intra-chromosomal structural variation events including inversions, translocations, aneuploidy (gain and loss), insertions, centromere abnormalities and complex events across the sample in additional cells from passage 21 (Figure 2).

#### Somatic variant annotation

An initial annotation of a publicly available HG008-T somatic variants called by DRAGEN v.4.2.4 was performed using ANNOVAR (v2020-06-08, https://annovar.openbioinformatics.org/en/latest/).^24–26^ DRAGEN calls were made from the Illumina-PCR-free-1 dataset. ANNOVAR was run using databases and annotation datasets: refGene, ClinVar (v20221231 and v20220320), Catalogue Of Somatic Mutations In Cancer (COSMIC) (v92, https://cancer.sanger.ac.uk/cosmic), gnomAD (v2.1), 1000 Genomes Project (2015 Aug), and the Exome Sequencing Project (ESP), as well as curation of genes commonly mutated in PDAC tumors.

#### HG008 ancestry analysis

We examined global genetic ancestry of the HG008 tumor and normal pancreatic samples, HG008-T and HG008-N-P (tissue) respectively, utilizing SNVs called by xAtlas from the Illumina-PCR-free-1 dataset. Ancestry was assessed by germline genetic similarity of these “target samples” to reference samples of known genographic origin from the 1000 Genomes Project (Phase 3).^27^ For computational efficiency, we limited our analysis to chromosome 21. We used PLINK^28^ to restrict our analysis to SNPs with minor allele frequency greater than or equal to 5 % in the reference panel. We then extracted these overlapping loci from the target sample Variant Call File (VCF) and then merged the datasets into a single VCF with bcftools.^29^ During the merging step, we assumed that entries absent from the target sample VCF at these sites were homozygous for the reference allele. We then performed principal component analysis (PCA) using PLINK and visualized genetic similarity on the first three principal component axes.

### Haplotype-specific copy number analysis

#### NIH-NCI

Haplotype-specific copy number analysis was performed using Wakhan (https://github.com/KolmogorovLab/Wakhan).^30^ Wakhan computes haplotype-specific coverage of the genome using phased heterozygous germline variants. Then it identifies boundaries of the copy number alteration (CNA) events by analysing SV breakpoints. Prior to running Wakhan, we phased the germline variants with PacBio long-read (Dataset ID: PB-HiFi-1) and Hi-C (Dataset ID: HiC-ILMN-2) data using HiPhase^31^ and HapCUT2.^32^

### STR genotyping

#### NIST

STR genotyping was performed for DNA isolated from the normal pancreatic and duodenal tissues and the HG008-T cells (batch 0823p23 and NIST passage 21) was performed using genomic DNA following a published protocol.^33^ Briefly, STR genotyping via capillary electrophoresis was conducted using PowerPlex Fusion 6C (Promega) and GlobalFiler (Thermo Fisher Scientific) according to the manufacturer’s protocols. The process targeted 1.0 ng of input DNA and utilized a 3500xL Genetic Analyzer with a 36 cm capillary array and POP-4 polymer (Thermo Fisher Scientific, Cat# A26070). Sample injection was performed at 1.2 kV for 15 seconds.

GeneMapper IDX v1.6 (Thermo Fisher Scientific) was employed for data interpretation, using the manufacturer’s provided bins and panels. Alleles were called using an analytical threshold set at 50 RFU. STR typing is provided in Table S4, see Supplementary Information document.

#### Data Records

All sequencing data presented are publicly accessible in the SRA database (SRP047086 under NCBI BioProject PRJNA200694).^8^ Bionano Optical Mapping data is provided as supplementary data on NCBI Supplementary Files (SUPPF_0000005645 under NCBI BioProject PRJNA200694). In addition to the G-banding and dGH karyograms shown in Figure 2, full G-banding cytogenetic analysis reports are on figshare.^34,35^

To aid in accessing the data for the various datasets presented, we provide a manifest (Table S1, see supplementary xlsx file) of all files with filenames, md5 checksums, and SRA associated identifiers. Full dataset descriptions are included in Table S1, see Supplementary Information document, and includes measurement technology, institution, sample source, sample(s) measured, e.g. cell line batch/passage and associated file types.

### Technical Validation

#### The tumor cell line has many characteristics of a typical tumor

To get a general overview of the aneuploidy and rearrangements in the tumor cell line, we performed standard karyotyping for MGH passage 31 (2022) and NIST passage 18 (2024), as well as karyotyping using Directional Genomic Hybridization (dGH) for NIST passage 21 (2024) of the tumor cell line; Dataset IDs karyotyping-1, karyotyping-2 and dGH-1, respectively. The karyotyping results indicate a subpopulation of the cell line has undergone whole genome doubling (Figure 2). Whole genome doubling generally is difficult to detect with bulk WGS, but is a common result of the *TP53* loss discussed below.^36^

To give an overview of copy number alterations in HG008-T, Figure 3 depicts the coverage by reads from each haplotype. The cell line appears to have many large deletions that cause loss of heterozygosity, with fewer large duplications. However, whole genome doubling in some cells results in four copies of regions without deletions and two copies of regions with deletions.

**Figure 3.**
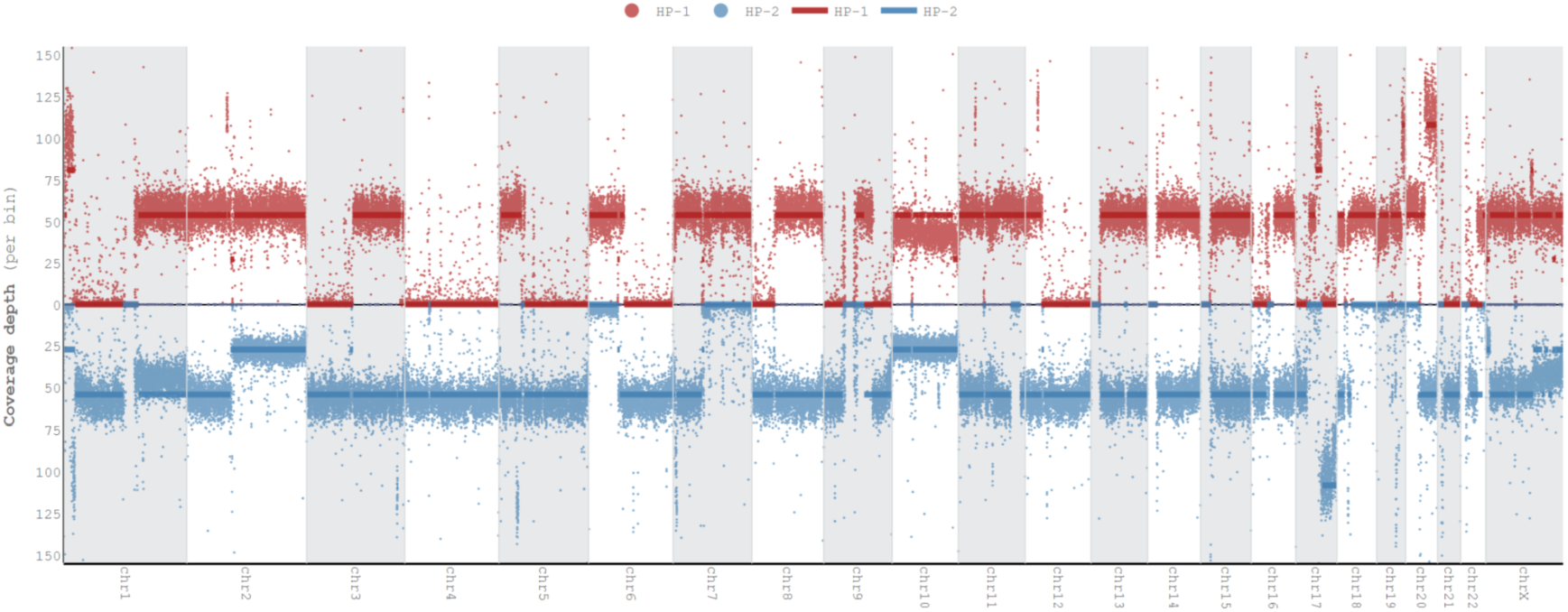
Haplotype Specific Coverage Plot. Plot shows coverage of haplotype 1 (HP-1, red) and haplotype 2 (HP-2, blue) for all chromosomes using HG008-T HiFi reads (Dataset ID: PB-HiFi-1) visualized using Wakhan (https://github.com/KolmogorovLab/Wakhan).

We performed an initial annotation of somatic variants^24^ and focused on somatic variants in genes commonly mutated in PDAC tumors. The cell line contains the *KRAS* [c.35G>T;p.G12V] (COSMIC ID: COSV55497419, and Pathogenic in ClinVar (https://www.ncbi.nlm.nih.gov/clinvar/variation/12583/) gain of function mutation common in PDAC^37^, as well as a likely duplication of the copy of the gene containing this mutation. In addition, there is a 20 kbp deletion of part of the *p16*/*CDKNA2* gene, and the other haplotype was lost due to a very large deletion. Other small somatic mutations that may have a functional impact on genes associated with cancer include the missense variant K132T [c.395A>C;p.Lys132Thr] in the remaining copy of the tumor suppressor gene *TP53* (COSV52759171, and likely pathogenic in ClinVar https://www.ncbi.nlm.nih.gov/clinvar/RCV000419605/),^38^ and a frameshift variant in the tumor suppressor gene *SMAD4* [c.153dup;p.Asp52ArgfsT], a variant not in COSMIC but in a gene associated with resistance to therapy.^39^ Therefore, HG008-T has somatic mutations in the four most commonly mutated genes in TCGA’s analysis of 150 PDAC tumors.^40^ Additional stopgain somatic variants in HG008-T and COSMIC include a SNV variant in *GLP2R* [c.1030C>T, p.R344*] (COSV52327429), and a SNV variant in *INSYN1* [c.457C>T, p.R153*] (COSV65838057). Note that these are examples of potentially important somatic variants in HG008-T, all of which appear to be in all tumor cells and not in normal cells, but are not a comprehensive list.

Also, it is possible that some of these may not have been in all cells in the primary tumor or may be new variants that occurred during cell line development. In addition, we compared the large-scale copy number alterations observed in HG008-T to recently published signatures of somatic copy alterations in TCGA samples,^41^ and the locations and prevalence of large deletions are similar to the chromosomal-scale loss of heterozygosity (cLOH) signature. The large-scale deletions followed by whole genome doubling in some cells is also consistent with the known progression of *TP53*-deficient tumors.^36^

### HG008 genetic ancestry analysis

To assess genetic ancestry of the HG008 individual, we performed principal component analysis (PCA) using PLINK^28^ and visualized genetic similarity on the first three principal component axes (Figure 4). The tumor and normal sample pair exhibited high genetic similarity on these axes, supporting their common germline origin, while also exhibiting the greatest genetic similarity to 1000 Genomes reference samples^27^ from the European superpopulation.

**Figure 4.**
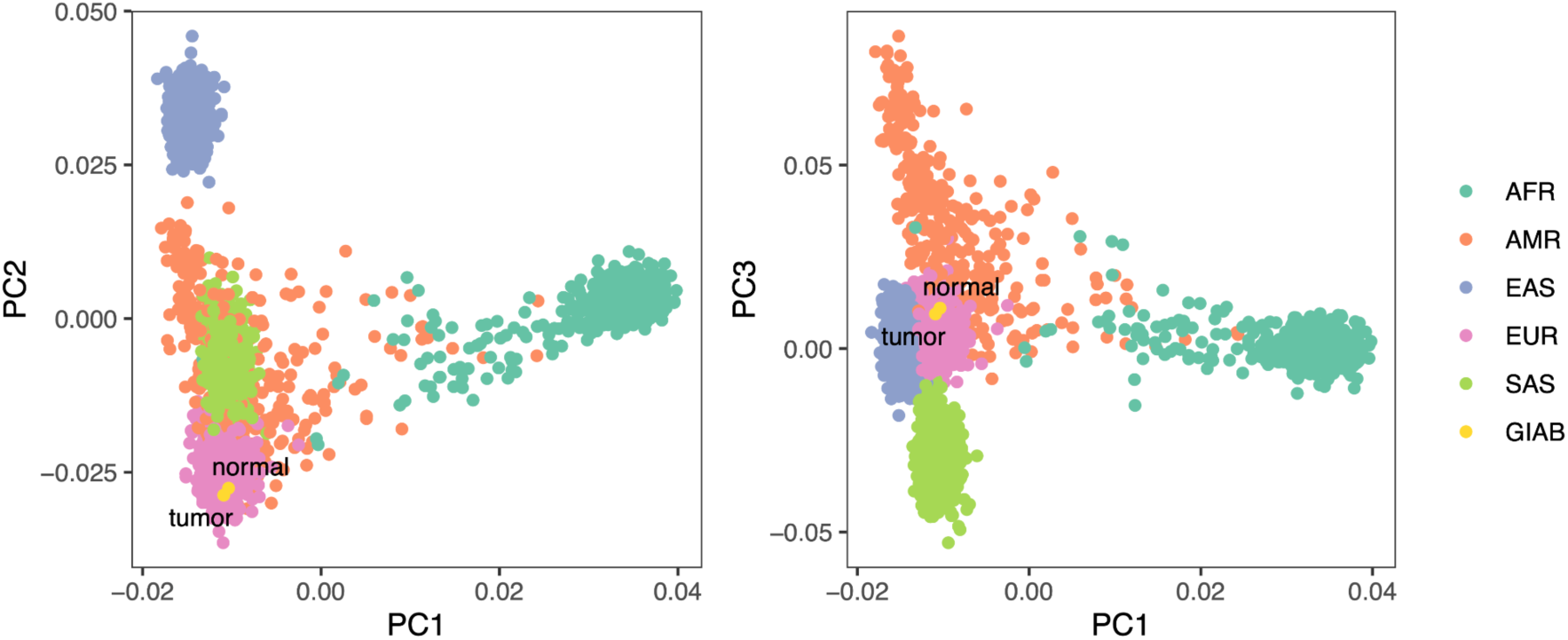
Ancestry Principal Component Analysis (PCA). PCA plot for HG008 using continental super populations of the 1000 Genomes reference samples. The PCA uses PLINK to visualize clusters of genetic similarity, and we show the first three principal component axes, which distinguish the super populations. Yellow points indicate both the normal and tumor cells’ variants from the HG008 individual make them most similar to individuals with European ancestry based on the first three principal components.

### PCR-free whole genome sequencing validation

To validate the quality of the sequencing data, we generated common sequencing QC metrics with public tools before and after mapping reads to the GRCh38-GIABv3 reference genome. This reference derives from the GRCh38 accession GCA_000001405.29, but without ALT loci, with false duplications masked, and with decoy sequences from T2T-CHM13v2.0 (accession GCA_009914755.4).^42^ We leveraged the GA4GH WGS quality control standard reference implementation (https://www.ga4gh.org/product/wgs-quality-control-standards/) for bulk WGS short-read data. When appropriate comparable QC tools were used for bulk WGS long-read data. More information on specific tools and metrics generated for validation of short-read and long-read data can be found in Tables S2 (tools) and S3 (metrics) in the Supplementary Information document. Overall, the QC metrics and data characteristics were consistent with expectations and the data were of sufficient quality for downstream analysis and use in material characterization. Additionally, the datasets have already been used for multiple downstream analyses, e.g. variant calling and genome assembly, further validating the data quality.

Because the tumor cell line is aneuploid, we also estimate the mean coverage of apparently diploid and haploid regions, as well as number of reads per tumor chromosomal copy (NRPCC).^41^ For these estimations, we use the mean coverage of chromosome 4 as the haploid coverage because it appears to be haploid in most or all cells, as was observed in the single cell data (Dataset ID: sc-ILMN-1). We calculate the diploid coverage as two times the haploid coverage and NRPCC as half the haploid coverage. The diploid and haploid coverages are estimated for cells without whole genome doubling, whereas NRPCC is the coverage of each copy of the chromosome for cells with whole genome doubling.

A summary of metrics generated by the GA4GH quality control pipeline for short-read sequencing technologies Ultima, Onso, Illumina, and Element are presented in Table 3. The short-read metrics show the read length and insert size mean and standard deviations were consistent with expected values for the library preparation methods used.

For long-reads, Table 4 has similar metrics for coverage but has some different metrics due to differences in QC tools used for long versus short-reads. For long-read technologies including ONT-std, ONT-UL, and Pacbio-Revio, QC tools such as Cramino and NanoPlot provide long-read specific metrics for validation (Table S2 (tools) and Table S3 (metrics), see Supplementary Information document). PycoQC metrics for the ONT datasets were generally consistent with expectations, with one exception. The first flowcell run for the ONT-std-1 dataset showed slightly lower read quality, Q18 compared to Q21 for subsequent flowcells measuring the HG008-N-P library, as reported by pycoQC. This is likely a result of flowcell variability and the flowcell being overloaded for the initial run, approximately 1.5X higher. For the two subsequent flowcells using the HG008-N-P library, the DNA mass input was adjusted down.

Figure 5a shows the read length distributions weighted by read length, similar to the N50 metric, and shows that ONT reads (particularly ultra long ONT, ONT-UL) are substantially longer than HiFi for the tumor cell line. However, because extracting very long DNA from normal tissues is more challenging than from cell lines, most ONT reads from normal pancreatic and duodenal tissue are shorter than from the tumor cell line. Figure 5c depicts this information in a different manner to make it easier to see the coverage by reads longer than a certain length (e.g., there is 33x coverage by ONT-UL-1 reads longer than 100 kb). Figure 5b shows the coverage has a single peak for normal cells but has two peaks for tumor cells due to apparently haploid and diploid regions.

**Figure 5.**
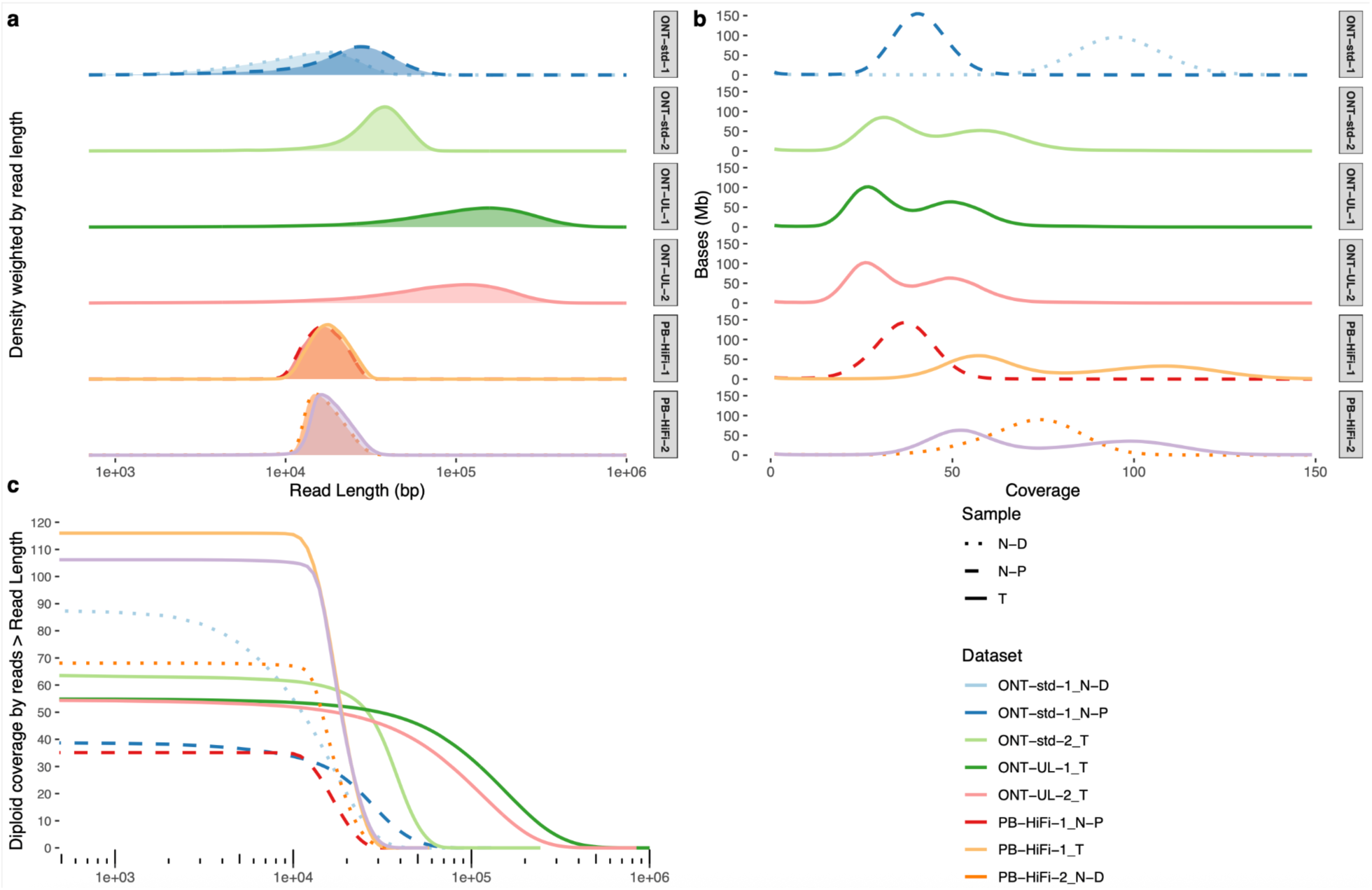
Long-read Coverage Plots. Read length and coverage distributions for long-read datasets **(a)** Mapped read length distribution weighted by aligned read length. **(b)** Coverage distribution showing number of genome positions (Mb) at each integer coverage. **(c)** Inverse cumulative distribution showing coverage of apparently diploid regions by aligned reads longer than the length on the x axis (this assumes no whole genome doubling).

#### Additional genome-scale measurement validation

For other datasets, QC metrics specific to each data type were generated and used to validate the datasets. The methods used for each validation are described in the associated Methods section above and in Table S2, see Supplementary Information document.

### Optical mapping validation

#### Bionano

All quality metrics for the Bionano-1 dataset passed manufacturer’s recommendations. The HG008-T sample had N50 of 322.82 kbp, labelling density (label/100 kbp) of 16.26, map rate of 92.2 % and effective coverage of 434.72X.

### BioSkryb single cell library validation

#### Illumina library validation

Single-cell libraries, from dataset sc-ILMN-1, were validated using the metrics reported by BioSkryb analytics BaseJumper platform using the BJ-DNA-QC pipeline (https://docs.basejumper.bioskryb.com/pipelines/secondary/bj-dna-qc/docs/). The BioSkryb BJ-DNA-QC pipeline metrics include genome coverage, percentage of mitochondrial contaminating reads, percentage of chimeric reads and assessing diversity of reads. A description of the BaseJumper reported metrics and their criteria for acceptance can be found in Table S3, see Supplementary Information document. Pre-seq counts are the estimation of the genomic coverage of a given library if it were to be run at the WGS level. In biological samples, pre-seq counts less than 3 billion can also represent biological genomic alterations (such as chromosomal losses). Samples with less than 3 billion pre-seq that meet other criteria are manually reviewed for copy number alteration to determine if biological features may be impacting this value. Because HG008-T contains substantial copy number alterations, particularly chromosome-scale losses, we are publishing data from cells that have pre-seq counts less than 3 billion.

Based on the metrics described above, a total of 98/120 libraries met the specifications by BioSkryb Genomics quality control. However, sequencing data for all 120 libraries is provided because the 22 libraries that did not pass BioSkryb QC might still be high quality libraries for cells with significant copy number alterations. Metrics for all 120 single-cell Illumina libraries are reported in Table S2 Bioskryb BJ-DNA QC (see supplementary xlsx file).

#### Ultima library validation

Ultima sequencing of the BioSkryb single cell libraries, presented in dataset sc-Ultima-1, was validated using the QC metrics reported from the UG100 sequencing run. Data is considered acceptable if the percentage of aligned reads is greater than 99 %, percentage of bases with a base quality of 20 is greater than 90 %, and the indel rate is less than 0.7 %. Library metrics were evaluated and 119/120 single cell libraries passed acceptance criteria. One library failed to meet acceptance criteria and was removed from the dataset. Metrics for the 119 single-cell Ultima libraries are reported in Table S2 Bioskryb Ultima QC (see supplementary xlsx file).

### HiC library validation

#### Phase Genomics

The HG008-T CytoTerra libraries, presented in datasets HiC-ILMN-1 and HiC-ILMN-3, were validated using the Phase Genomics hiC_QC.py script (https://github.com/phasegenomics/hic_qc). Both libraries met all Phase Genomics CytoTerra QC acceptance criteria as described below and in Table S3, see Supplementary Information document. The following describes the observed metrics for HiC-ILMN-1 and HiC-ILMN-3, respectively. High-quality read pairs are those having MAPQ greater than zero, are not PCR duplicates and map to different contigs or are greater than10 kb apart. The percentage of high-quality read pairs where both reads map to the same strand in the reference genome is 47.54 % and 36.56 %, well exceeding the minimum acceptable criteria of 15 %. The fraction of high-quality read pairs greater than 10 kb apart is 75.27 % and 43.51 %, exceeding minimum acceptable criteria of 20 %. The fraction of duplicate reads is 21.28% and 35.79 %, below the maximum acceptable criteria of 40 %. Finally, the frequency of pairs of reads mapping to different contigs in an assembly (intercontig read pairs) is 22.25 % and 14.15%, below the maximum acceptance criteria of 30%.

#### Arima Genomics

The HG008-T and HG008-N-D HiC libraries and resulting sequencing, presented in the HiC-ILMN-2 dataset, were found to meet criteria for acceptance. The percentage of single-end reads (%_Mapped_SE_reads) that can be mapped to the reference, out of the total number of single-end reads, is greater than 80 %. The percent of all unique total pairs that have both read-ends aligning to the same chromosome and have an insert size greater than or equal to 15 kb (percent long-cis (greater than 15 kb)) is greater than 30 % and the ratio of long-cis to Trans data (Lcis_trans_ratio) is greater than one. This ratio is the signal to noise ratio for translocation calling. HiC data QC statistics for the deeply sequenced Illumina data can be found in Table S2 Arima Deep QC (see supplementary xlsx file).

#### Dovetail Genomics

The HG008-T LinkPrep libraries and resulting sequencing, presented in the HiC-ILMN-4 dataset, were found to meet criteria for acceptance. Libraries are considered passing for QC metrics as follows; mapping rate greater than 50%, Duplicate rate less than 50 %, and the % Cis read pairs separated by than 1kb (% Long Range) has a value greater than 20 %. LinkPrep library QC statistics from Illumina sequencing can be found in Table S2 Dovetail QC (see supplementary xlsx file).

#### Usage Notes

The data presented herein were collected as part of the Genome in a Bottle (GIAB) Consortium’s efforts (https://www.nist.gov/programs-projects/cancer-genome-bottle). The data and alignments are publicly available on the GIAB FTP site hosted by the NIH-NCBI (https://ftp.ncbi.nlm.nih.gov/ReferenceSamples/giab/data_somatic/HG008/). Additionally, the sequencing data can be browsed through the 42basepairs platform, which provides access to both the GIAB FTP site (https://42basepairs.com/browse/web/giab/data_somatic/HG008/) and the GIAB S3 bucket (https://42basepairs.com/browse/s3/giab/data_somatic/HG008), allowing users to obtain S3 URLs. The 42basepairs platform enables high-level exploration of the data. Future data and analyses generated for the GIAB HG008 sample will be made publicly available on the GIAB FTP site as a resource for the community.

We expect these extensive data sets to be useful for methods development for somatic variant calling, personalized tumor-normal analysis, and tumor genome assembly. As GIAB is developing initial somatic benchmarks for batch 0823p23 of this tumor cell line, methods can be optimized and evaluated against these benchmarks. The sequencing coverages in this work are more than sufficient for identifying clonal somatic variants that are likely to be consistent between passages of the tumor cell line. In the long-term, larger batches of cells from the tumor cell line can be characterized for differences in somatic variants relative to batch 0823p23, enabling a more sustainable supply of DNA and cells. In addition, extensive data from two different normal tissues from the same individual can enable understanding of mosaicism. Long reads also contain methylation tags for epigenetic analyses. Future work will be needed to understand the genomic stability of the tumor cell line. Our initial data from different passages can be used to gain a preliminary understanding of stability, and we plan to add more data from different passages performed at different laboratories in the future.

As noted previously, we are working to deposit the tumor cell line in a public repository. To aid in cell line authentication of HG008-T cells deposited in cell repositories, we have provided the STR profile for the HG008-T cell line and normal tissues in Table S4, see Supplementary Information document. As additional data, benchmark, and cell line resources become available, information for these resources will be posted on the NIST Cancer Genome in a Bottle webpage (https://www.nist.gov/programs-projects/cancer-genome-bottle).

## Code Availability

Scripts used for validation and figures are located on GitHub at https://github.com/usnistgov/giab-HG008-data-paper.

## Acknowledgements

The work of H.D., S.N.J., D.M.M., M.G., H.M., S.B. and F.J.S from Baylor College of Medicine was performed with support from NIH S10 grant 1S10OD028587. The work of F.T-N., J.T. and C.X. were supported by the National Center for Biotechnology Information of the National Library of Medicine (NLM), National Institutes of Health. Sequencing and analysis performed by G.N., Z.S., C.R., and J.S. from NYGC were supported by the National Cancer Institute of the National Institutes of Health under Award Number U01CA253405. J.P., I.V., K.H.M., J.G., and B.M from UCSC were supported by the National Institutes of Health under Award Number U24HG011853. K.H.M. from UCSC was supported by a grant from NHGRI, R01HG011274. G.R. from Drexel was supported by grants from NSF (#1936791, #1919691 and #2107108). R.M from JHU was supported by grants from NIH (R35GM133747 and OT2OD034190). Establishment of the HG008-T cell line by A.L at MGH was supported by a grant from NIST, 60NANB20D166. Certain commercial equipment, instruments, or materials are identified to specify adequately experimental conditions or reported results. Such identification does not imply recommendation or endorsement by the National Institute of Standards and Technology, nor does it imply that the equipment, instruments, or materials identified are necessarily the best available for the purpose.

## Author contributions

NIST coordinated data acquisition and writing of the manuscript. J.M., V.P. and N.D.O. performed data management and QC, J.W. performed analysis, J.H.M., V.P. and J.M.Z. writing. K.D.C. initiated communications with MGH to establish tumor-normal paired cell lines. A.A.G. performed STR typing.

Additionally, J.M.Z. prepared experimental designs for measurements.

Z.H. and H-J.H. from NIST performed cell culturing of the HG008-T cell line.

A.S. and K.S. generated Arima HiC data and performed analysis.

For BCM, project management was performed by H.D. and S.N.J. H.D., S.N.J., D.M.M., M-C.G. and F.S. generated Illumina and PacBio Revio data, and S.B, L.F.P. and F.J.S. performed analysis.

A.R.H. and H-C.Y. generated Bionano Genomics data and performed analysis. H-C.Y. wrote methods.

For BioSkryb Genomics, A.R. and K.K. performed FACs sorting, K.K. and J.R. performed experimental amplification and QC sequencing and I.G. and V.W. performed analysis.

For Element Biosciences, M.S. and S.L. provided project management, M.S. and K.W. generated Element data and wrote methods and K.W. and B.R.L. performed analysis.

M.J. and S.K. from Northeastern University prepared samples and generated ONT data, M.J. performed QC, data management and wrote methods.

G.N., Z.S., C.R., and J.S. from NYGC generated Illumina data and G.N. and J.S. performed analysis.

For PacBio Revio, C.L., P.B., and I.J.M. generated data, A.M.W. performed analysis, and S.B.K. wrote methods. For PacBio Onso, A.A. generated data,

C.K. performed analysis, and C.K., Y.K., and M.W. wrote methods.

J.P., I.V., K.H.M., J.G., and B.M. from UCSC generated ONT data.

G.L.R. from Drexel University performed Illumina data analysis.

R.M. from JHU performed ancestry analysis and participated in writing methods.

E.B., E.S., K.S. and S.T. from Illumina, Inc. generated data and performed analysis.

L.C.H. from the NIH/NCI, Division of Cancer Epidemiology and Genetics prepared SNV annotations and variant interpretation and review of SNV function.

For Ultima Genomics, N.I. generated UG100 data, D.L., H.B., I.S. and G.K. performed analysis and D.L. provided project management and wrote methods.

S.E. from Phase Genomics generated data and S.E. and M.W. performed analysis.

For KROMATID, M.V. generated data, E.C. and M.V. performed QC, E.C., S.G., and M.V. performed analysis, G.H. provided project management and E.C. generated the experimental approach for data generation and wrote methods.

M.K., T.A., A.K. and A.B. from the NIH/NCI Cancer Data Science Laboratory, performed Haplotype-specific copy number analysis.

F.T-N., J.T. and C.X. from the NIH/NCBI performed data management and QC. C.X. was lead data manager for NCBI GIAB ftp and aws s3.

J.P., J.W.H., K.R. and C.E.M. from Weill Cornell Medicine generated data, K.R. and C.E.M performed analysis, J.P. handled project logistics, J.W.H. performed project management and C.E.M also provided experimental design.

E.M.M. from Cantata Bio performed data analysis and project management (Dovetail LinkPrep). M.S.B. performed Library Prep and data generation (Dovetail LinkPrep). J.Z.S. performed Data analysis (Dovetail LinkPrep).

A.S.L. from MGH established the HG008-T cell line and distributed tumor and normal samples for measurements.

