## Supplementary Information Document for "Development and extensive sequencing of a broadly-consented Genome in a Bottle matched tumor-normal pair"

### Supplementary Information Tables

1. Table S1 Data records
2. Table S2 Validation tools
3. Table S3 QC metrics descriptions
4. Table S4 HG008 STR typing

| Measurement Technology | Measurement Institution | Dataset ID | Sample Source | Sample Material Received | File Type(s) | Number of files | Data Access |
|---|---|---|---|---|---|---|---|
| Illumina NovaSeq 6000 PCR-Free WGS | Baylor College of Medicine, Human Genome Sequencing Center | ILMN-PCR-free-1 | normal pancreatic tissue | tissue | paired-end reads (R1/R2) in fastq.gz format | x2 (fastq.gz, R1/R2) | SRA |
| | | | PDAC tumor cell line | initial growth 2022, p36 cells | | x2 (fastq.gz, R1/R2) | |
| | | ILMN-PCR-free-2 | normal duodenal tissue | tissue | | x2 (fastq.gz, R1/R2) | |
| | | | PDAC tumor cell line | batch 0823p23 cells | | x8 (fastq.gz, R1/R2) | |
| | New York Genome Center | ILMN-PCR-free-3 | normal duodenal tissue | isolated DNA | | x24 (fastq.gz, R1/R2) | |
| | | | PDAC tumor cell line | batch 0823p23 cells | | x24 (fastq.gz, R1/R2) | |
| Element Aviti | Element Biosciences | Element-short-3 | normal duodenal tissue | isolated DNA | paired-end reads (R1/R2) in fastq.gz format | x2 (fastq.gz, R1/R2) | |
| | | | normal pancreatic tissue | isolated DNA | | x2 (fastq.gz, R1/R2) | |
| | | | PDAC tumor cell line | batch 0823p23 isolated DNA | | x2 (fastq.gz, R1/R2) | |
| | | Element-long-1 | normal duodenal tissue | isolated DNA | | x2 (fastq.gz, R1/R2) | |
| | | | normal pancreatic tissue | isolated DNA | | x2 (fastq.gz, R1/R2) | |
| | | | PDAC tumor cell line | batch 0823p23 isolated DNA | | x2 (fastq.gz, R1/R2) | |
| PacBio Onso | PacBio | PB-Onso-1 | normal duodenal tissue | isolated DNA | paired-end reads (R1/R2) in fastq.gz format | x2 (fastq.gz, R1/R2) | |
| | | | PDAC tumor cell line | batch 0823p23 isolated DNA | | x2 (fastq.gz, R1/R2) | |
| Ultima UG100 | Ultima Genomics | Ultima-bulk-1 | normal duodenal tissue | isolated DNA | aligned reads in .cram format with associated .cram.crai index | x2 (.cram/.crai) | |
| | | | PDAC tumor cell line | batch 0823p23 isolated DNA | | x2 (.cram/.crai) | |
| | | Ultima-ppmSeq-2 | normal duodenal tissue | isolated DNA | | x2 (.cram/.crai) | |
| | | | PDAC tumor cell line | batch 0823p23 isolated DNA | | x2 (.cram/.crai) | |
| HiC-Illumina | Phase Genomics | HiC-ILMN-1 | PDAC tumor cell line | initial growth 2022, p38 viable cells | paired-end reads (R1/R2) in fastq.gz format | x4 (fastq.gz, R1/R2) | |
| | Arima Genomics - Baylor College of Medicine | HiC-ILMN-2 | normal duodenal tissue | tissue | | x2 (fastq.gz, R1/R2) | |
| | | | PDAC tumor cell line | batch 0823p23 cells | | x2 (fastq.gz, R1/R2) | |
| | Phase Genomics | HiC-ILMN-3 | PDAC tumor cell line | 2024 NIST p21 cells | | x2 (fastq.gz, R1/R2) | |
| | Dovetail Genomics | HiC-ILMN-4 | PDAC tumor cell line | 2024 NIST p21 cells | | x8 (fastq.gz, R1/R2) | |
| PacBio Revio (HiFi) | PacBio | PB-HiFi-1 | normal pancreatic tissue | isolated DNA | raw reads in unaligned .bam format | x1 (.bam) | |
| | | | PDAC tumor cell line | batch 0823p23 isolated DNA | | x2 (.bam) | |
| | Baylor College of Medicine, Human Genome Sequencing Center | PB-HiFi-2 | normal duodenal tissue | tissue | | x2 (.bam) | |
| | | | PDAC tumor cell line | batch 0823p23 cells | | x2 (.bam) | |
| Oxford Nanopore Technologies (ONT) PromethION (standard - simplex) | Northeastern University, Genome Technology Lab | ONT-std-1 | normal duodenal tissue | tissue | raw reads in unaligned .bam format | x4 (.bam) | |
| | | | normal pancreatic tissue | tissue | | x3 (.bam) | |
| | University of CA Santa Cruz Genomics Institute | ONT-std-2 | PDAC tumor cell line | batch 0823p23 cells | | x1 (.bam) | |
| Oxford Nanopore Technologies (ONT) PromethION (Ultra Long) | University of CA Santa Cruz, Genomics Institute | ONT-UL-1 | PDAC tumor cell line | batch 0823p23 cells | | x3 (.bam) | |
| | Northeastern University, Genome Technology Lab | ONT-UL-2 | PDAC tumor cell line | batch 0823p23 cells | | x1 (.bam) | |
| Single-Cell (BioSkryb ResolveDNA) WGS Illumina | BioSkryb Genomics | sc-ILMN-1 | PDAC tumor cell line | batch 0823p23 cells | paired-end reads (R1/R2) in fastq.gz format | x240 (.fastq.gz, R1/R2) | |
| Single-Cell (BioSkryb ResolveDNA) WGS Ultima | BioSkryb - Ultima Genomics | sc-Ultima-1 | PDAC tumor cell line | Converted sc-ILMN-1 libraries to Ultima libraries | aligned (GRCh38) reads in . cram format with associated . cram.crai index | x238 (.cram./.crai) | |
| Saphyr Optical Mapping | Bionano Genomics | Bionano-1 | PDAC tumor cell line | batch 0823p23 cells | raw data view of molecule and label information in .bnx format and description of SVs detected by two genome maps in .smap format | x3 (.bnx/.smap/.cmap) | NCBI supplementary |
| Karyotyping | KaryoLogic | karyotpying-1 | PDAC tumor cell line | initial growth 2022, p31 viable cells | karyogram provided in cytogenetic analysis report (.pdf) | x2 (.pdf/.xlsx) | figshare |
| | | karyotpying-2 | PDAC tumor cell line | 2024 NIST p18 cells | | x1 (.pdf) | |
| Directional Genomic Hybridization (dGH) KROMASURE Screen | KROMATID | dGH-1 | PDAC tumor cell line | 2024 NIST p21 cells | karyogram provided as part of Figure 2 | NA | karyogram provided as part of Figure 2 |

### Table S1 Dataset descriptions

Summary of HG008 datasets, including measurement institution, file types, and data access information. The sample material received denotes the starting material received by the measurement institution; cells, tissue or isolated genomic DNA. Tumor samples derive from either a large, homogeneous batch of cells distributed in 2023 (0823p23) by the Liss lab at MGH, from a series of 2022 Liss lab passages starting with the p13 stockthat batch 0823p23 was grown from or from NIST passages in 2024 started from Liss lab p13 cryopreserved cells.

| Dataset ID | Mapping Tools | | Reference Genome | Validation Tools | |
|---|---|---|---|---|---|
| ILMN-PCR-free-1 | BWA-MEM | BWA: https://github.com/lh3/bwa | GRCh38-GIABv3: https://ftp-trace.ncbi.nlm.nih.gov/giab/ftp/release/references/GRCh38/GRCh38_GIABv3_no_alt_analysis_set_maskedGRC_decoys_MAP2K3_KMT2C_KCNJ18.fasta.gz | FastQC<br>GA4GH QC pipeline<br>Mosdepth (run as part of snakemake workflow) | FastQC: https://github.com/s-andrews/FastQC<br>GA4GH Pipeline: https://github.com/c-BIG/NPM-sample-qc/tree/v0.11.0/<br>Mosdepth Snakemake wrapper: https://snakemake-wrappers.readthedocs.io/en/stable/wrappers/mosdepth.html |
| ILMN-PCR-free-2 | | | | | |
| Element-short-3 | | | | | |
| Element-long-1 | | | | | |
| PB-Onso-1 | | | | | |
| ILMN-PCR-free-3 | | NYGC somatic pipeline v7: https://bitbucket.nygenome.org/projects/WDL/repos/somatic_dna_wdl/browse/README_pipeline.md?at=refs%2Fheads%2Fgiab#deliverables | | | |
| Ultima-bulk-1 | Optimized version of BWA (UA) for Ultima data | Ultima Aligner: https://hub.docker.com/r/ultimagenomics/alignment | | GA4GH QC pipeline<br>Mosdepth (run as part of snakemake workflow) | GA4GH Pipeline: https://github.com/c-BIG/NPM-sample-qc/tree/v0.11.0/<br>Mosdepth Snakemake wrapper: https://snakemake-wrappers.readthedocs.io/en/stable/wrappers/mosdepth.html |
| HiC-ILMN-1 | BWA-MEM (as part of the HiC alignment and QC protocol) | HiC alignment and QC: https://phasegenomics.github.io/2019/09/19/hic-alignment-and-qc.html | GRCh38.p13: https://www.ncbi.nlm.nih.gov/datasets/genome/GCF_000001405.39/ | hic_qc (as part of the HiC alignment and QC protocol) | HiC QC script: https://github.com/phasegenomics/hic_qc |
| HiC-ILMN-3 | | | | | |
| HiC-ILMN-2 | HiCUP (as part of the Arima SV pipeline) | Arima SV pipeline: https://arimagenomics.com/wp-content/files/Bioinformatics-User-Guide-Arima-Structural-Variant-Pipeline.pdf | hg38: no path provided | QC part of the Arima SV pipeline | Arima SV pipeline: https://arimagenomics.com/wp-content/files/Bioinformatics-User-Guide-Arima-Structural-Variant-Pipeline.pdf |
| HiC-ILMN-4 | BWA-MEM (as part of the Alignment and Proximity-Ligation QC) | Alignment and Proximity-Ligation QC: https://varilink.readthedocs.io/en/latest/library_qc.html | hg38: https://hgdownload.soe.ucsc.edu/goldenPath/hg38/bigZips/hg38.fa.gz | QC output from pairtools (as part of the Alignment and Proximity-Ligation QC) | pairtools: https://github.com/open2c/pairtools |
| PB-HiFi-1 | pbmm2 (as part of PacBio WDL germline (v1.1.1) or somatic (v0.7) pipelines) | pbmm2: https://github.com/PacificBiosciences/pbmm2<br>PacBio germline pipeline v1.1.1: https://github.com/PacificBiosciences/HiFi-human-WGS-WDLPacBio<br>somatic pipeline v0.7: https://github.com/PacificBiosciences/HiFi-Somatic-WDL | GRCh38-GIABv3: https://ftp-trace.ncbi.nlm.nih.gov/giab/ftp/release/references/GRCh38/GRCh38_GIABv3_no_alt_analysis_set_maskedGRC_decoys_MAP2K3_KMT2C_KCNJ18.fasta.gz | Cramino<br>Nanoplot<br>Mosdepth<br>Samtools stats | Cramino: https://github.com/wdecoster/cramino<br>Nanoplot: https://github.com/wdecoster/NanoPlot<br>Mosdepth Snakemake wrapper: https://snakemake-wrappers.readthedocs.io/en/stable/wrappers/mosdepth.html<br>Samtools: https://github.com/samtools/samtools |
| PB-HiFi-2 | | | | | |
| ONT-std-1 | Samtools<br>Minimap2 | Samtools: https://github.com/samtools/samtools<br>minimap2: https://github.com/lh3/minimap2 | | PycoQC<br>Cramino<br>Nanoplot<br>Mosdepth<br>Samtools stats | pycoQC: https://github.com/a-slide/pycoQC<br>Cramino: https://github.com/wdecoster/cramino<br>Nanoplot: https://github.com/wdecoster/NanoPlot<br>Mosdepth Snakemake wrapper: https://snakemake-wrappers.readthedocs.io/en/stable/wrappers/mosdepth.html<br>Samtools: https://github.com/samtools/samtools |
| ONT-std-2 | | | | | |
| ONT-UL-1 | | | | | |
| ONT-UL-2 | | | | | |
| sc-ILMN-1 | Sention BWA-MEM (as part of BJ-DNA-QC pipeline) | BJ-DNA-QC pipeline: https://docs.basejumper.bioskryb.com/pipelines/secondary/bj-dna-qc/1.8.2/docs/ | GRCh38: * gs://gcp-public-data--broad-references/hg38/v0/Homo_sapiens_assembly38.fasta | validation as part of the BJ-DNA-QC pipeline | BJ-DNA-QC pipeline: https://docs.basejumper.bioskryb.com/pipelines/secondary/bj-dna-qc/1.8.2/docs/ |
| sc-Ultima-1 | Optimized version of BWA (UA) for Ultima data | Ultima Aligner: https://hub.docker.com/r/ultimagenomics/alignment | GRCh38: * gs://gcp-public-data--broad-references/hg38/v0/Homo_sapiens_assembly38.fasta | Ultima custom UG100 tool | Ultima Sorter: https://hub.docker.com/r/ultimagenomics/sorter |
| Ultima-ppmSeq-1 | Optimized version of BWA (UA) for Ultima data | Trim, align and sort using ppmSeqPreprocess workflow: https://github.com/Ultimagen/healthomics-workflows/tree/0c827f223b47d84736cea7626af6ddb8ca0a90a0/workflows/ppmSeq_preprocess | GRCh38: * gs://gcp-public-data--broad-references/hg38/v0/Homo_sapiens_assembly38.fasta | GA4GH QC pipeline | GA4GH pipeline: https://github.com/c-BIG/NPM-sample-qc/tree/v0.11.0/ |
| Bionano-1 | Bionano Rare Variant Analysis pipeline (included in Bionano Access software) | Bionano proprietary tool | GRCh38: https://ftp.ncbi.nlm.nih.gov/genomes/all/GCA/000/001/405/GCA_000001405.15_GRCh38/seqs_for_alignment_pipelines.ucsc_ids/GCA_000001405.15_GRCh38_no_alt_analysis_set.fna.gz | Bionano Access version 1.8 | Bionano proprietary tool |

**Table S2 Validation Tools**

Summary of tools used for alignment and validation.

*Information for accessing and use of files from the Broad Resoruce bundle can be found at https://gatk.broadinstitute.org/hc/en-us/articles/360035890811-Resource-bundle

| Tool | Metric | Description |
|---|---|---|
| Cramino | number of alignments | Total number of reads aligned to the reference genome. |
| | percent from total reads | Percentage of reads successfully aligned. |
| | yield (Gb) | The total amount of sequencing data generated, measured in gigabases (Gb). |
| | mean coverage | The average number of times each base in the genome is read during sequencing. |
| | N50 | Half of the sequenced bases are in reads longer than the N50. |
| | median identity to GRCh38 | Median fraction (%) of read bases matching the GRCh38 reference genome. |
| Samtools stats | samtools error rate (%) | The proportion (%) of bases in the reads that do not match the reference. The output of samtools stats is commonly referred to as the error rate. The samtools error rate measures substitution and indel differences between the reads and the reference, including differences resulting from true variants. |
| | average length (bp) | The average length of the reads in the dataset. |
| | avg. insert size (bp) | The average length of the DNA fragments that are sequenced. The output of the samtools stats is commonly referred to as the insert size average. |
| | insert size SD | The variability in the insert sizes. The output of the samtools stats is commonly referred to as the insert size standard deviation. |
| | properly paired (%) | The percentage of read pairs that are aligned with the expected pairing. The output of the samtools stats is commonly referred to as the percentage of properly paired reads. |
| | mapped reads (%) | The percentage of reads mapping to the reference genome. This is calculated using outputs from samtools stats ("reads mapped" / "raw total sequences" x 100). |
| | bases mapped (%) | The percentage of bases, ignoring clipping, that map to the reference genome. This is calculated using outputs from samtools stats, (bases mapped (cigar))/(total length) x 100). |
| GA4GH mosdepth | mean autosome coverage | The mean sequencing coverage derived from high-quality, non duplicated reads, primary alignments in the autosome non-gap regions of the reference used. |
| Mosdepth | haploid mean coverage | Estimated coverage of haploid regions assuming no whole genome doubling, derived from the mean sequencing coverage derived from high-quality, non duplicated reads, primary alignments in chromosome 4 for HG008-T, which is haploid. |
| | diploid mean coverage | Estimated coverage of diploid regions assuming no whole genome doubling, calculated as double the haploid mean coverage. |
| | NRPCC | Estimated coverage per haplotype assuming whole genome doubling has occurred. |
| BioSkryb BJ-DNA-QC pipeline metrics | Pre-Seq Count | Estimation of the genomic coverage of a given library if it were to be run at the WGS level. Traditionally, BioSkryb Genomics has utilized the NIST cell line NA12878,in this cell line, which has no major genomic alterations, pre-seq counts of high-quality cells are typically greater than 3,000,000,000. |
| | % ChrM | Percentage of mitochondrial reads in the sequencing, defined as <2% for passing cells. |
| | % Chimera | Percentage of chimeric reads seen at the QC Sequencing level, defined as <20% for passing cells. |
| | Gini coefficient | The Gini coefficient assess assay and sequencing performance by measuring the diversity of reads sequenced, diversity of reads is reflective of homogeneity of coverage, the higher the homogeneity of coverage the greater the diversity of reads sequences. This is also typically associated with the ability to assess copy number variation at the QC sequencing level. Passing cells are below 0.1 |
| Phase Genomics CytoTerra QC metrics | High-Quality | Refers to read pairs having MAPQ>0, are not PCR duplicates and map to different contigs or are >10kb apart. |
| | Same strand high quality read pairs | This statistic measures the percentage of high quality read pairs where both reads map to the same strand in the reference genome. This is indicative that the read is the result of a proximity ligation event which changes the orientation of the sequences relative to each other. Doubling this value gives an estimate of the total percentage of proximity ligation junctions present in the library. (15% minimum value acceptable) |
| | Fraction of high quality read pairs >10 kb apart | CytoTerra library success is dependent on the fraction of reads that contain long-range contact information. This stat measures the percentage of high quality read pairs that map >10 kb apart in the reference genome. (20% minimum acceptable value) |
| | Duplicate Reads | This measures the rate of PCR duplicate fragments present in the library and fits a saturation model to extrapolate the duplication rate at 100M read pairs. This is a critical measure of the complexity of a library. (40% maximum acceptable value). |
| | Intercontig read pairs | This measures the frequency which pairs of reads map to different contigs in an assembly. Since in the human genome, contigs are (nearly) complete chromosomes, intercontig read pairs is an estimate of spurious proximity ligation events. There are exceptions to this threshold in highly rearranged genomes since you then expect interchromosomal read pairs due to translocations. (30% maximum acceptable value) |

**Table S3 Validation metrics descriptions**

Overview of tools and description of their output metrics presented as part of validation for the HG008 datasets.

| Normal Duodenal Tissue | |
|---|---|
| AMEL | X |
| D3S1358 | 14, 15 |
| D1S1656 | 16, 17.3 |
| D2S441 | 11 |
| D10S1248 | 14 |
| D13S317 | 11, 12 |
| Penta E | 12, 18 |
| D16S539 | 11 |
| D18S51 | 10, 15 |
| D2S1338 | 17, 18 |
| CSF1PO | 10, 11 |
| Penta D | 12 |
| TH01 | 8, 9.3 |
| vWA | 14, 18 |
| D21S11 | 28, 29 |
| D7S820 | 7, 8 |
| D5S818 | 11 |
| TPOX | 10, 11 |
| D8S1179 | 10, 13 |
| D12S391 | 17, 18 |
| D19S433 | 12, 13 |
| SE33 | 18, 23 |
| D22S1045 | 11, 15 |
| FGA | 21, 25 |

| Normal Pancreatic Tissue | |
|---|---|
| AMEL | X |
| D3S1358 | 14, 15 |
| D1S1656 | 16, 17.3 |
| D2S441 | 11 |
| D10S1248 | 14 |
| D13S317 | 11, 12 |
| Penta E | 12, 18 |
| D16S539 | 11 |
| D18S51 | 10, 15 |
| D2S1338 | 17, 18 |
| CSF1PO | 10, 11 |
| Penta D | 12 |
| TH01 | 8, 9.3 |
| vWA | 14, 18 |
| D21S11 | 28, 29 |
| D7S820 | 7, 8 |
| D5S818 | 11 |
| TPOX | 10, 11 |
| D8S1179 | 10, 13 |
| D12S391 | 17, 18 |
| D19S433 | 12, 13 |
| SE33 | 18, 23 |
| D22S1045 | 11, 15 |
| FGA | 21, 25 |

| HG008-T cell line (batch 0823p23) | |
|---|---|
| AMEL | X |
| D3S1358 | 15 |
| D1S1656 | 16, 17.3 |
| D2S441 | 11 |
| D10S1248 | 14 |
| D13S317 | 11, 12 |
| Penta E | 12, 18 |
| D16S539 | 11 |
| D18S51 | 10 |
| D2S1338 | 17, 18 |
| CSF1PO | 11 |
| Penta D | 12 |
| TH01 | 8, 9.3 |
| vWA | 14, 18 |
| D21S11 | 28 |
| D7S820 | 8 |
| D5S818 | 11 |
| TPOX | 10, 11 |
| D8S1179 | 10, 13 |
| D12S391 | 17, 18 |
| D19S433 | 12 |
| SE33 | 23 |
| D22S1045 | 15 |
| FGA | 25 |

| HG008-T cell line (NIST passage 21) | |
|---|---|
| AMEL | X |
| D3S1358 | 15 |
| D1S1656 | 16, 17.3 |
| D2S441 | 11 |
| D10S1248 | 14 |
| D13S317 | 11, 12 |
| Penta E | 12, 18 |
| D16S539 | 11 |
| D18S51 | 10 |
| D2S1338 | 17, 18 |
| CSF1PO | 11 |
| Penta D | 12 |
| TH01 | 8, 9.3 |
| vWA | 14, 18 |
| D21S11 | 28 |
| D7S820 | 8 |
| D5S818 | 11 |
| TPOX | 10, 11 |
| D8S1179 | 10, 13 |
| D12S391 | 17, 18 |
| D19S433 | 12 |
| SE33 | 23 |
| D22S1045 | 15 |
| FGA | 25 |

**Table S4 STR genotyping**

STR genotyping was conducted on DNA isolated from normal pancreatic and duodenal tissues, as well as the HG008-T cells (batch 0823p23 and NIST passage 21). The resulting allelic profiles for each marker are presented in the accompanying table. Notably, some markers in the HG008-T samples exhibited allelic loss due to large deletions previously identified in other analyses, as discussed in the Technical Validation section.
