## Supplementary File 1 for "Development and extensive sequencing of a broadly-consented Genome in a Bottle matched tumor-normal pair"

### UG ppmSeq™ Genomic DNA Library Preparation Protocol

Optimized with NEBNext UltraShear™, NEBNext® Ultra™ II End Repair/dA-Tailing Module, NEBNext® Ultra™ II Ligation Module

Rev. 01

UG ppmSeq™ Genomic DNA Library Preparation Protocol
D1000993 Rev. 01 | 30 September 2024

© 2024, Ultima Genomics, Inc. All rights reserved.

UG 100™, ppmSeq™, ULTIMA GENOMICS®, and the Ultima Genomics logo are trademarks of Ultima Genomics, Inc. Other product names and trademarks are the property of their respective owners.

### 1. Revision History

| Revision | Release Date | Description of Changes |
|---|---|---|
| 01 | 10/30/2024 | Initial release |

### 2. Document Conventions

The following conventions are used in this document.

#### Icons and Text Conventions

| Icon | Description |
|---|---|
| | **NOTE**<br>Alerts you of important information about a system feature or process. |
| | **CAUTION!**<br>Alerts you of a hazardous situation that could result in potential loss of data or damage to the system. |
| | **WARNING!**<br>Alerts you of a hazardous situation that could result in potential personal injury. |
| | **SAFE STOPPING POINT!**<br>Indicates a safe stopping point in a procedure. |
| Hyperlink | Hyperlinks are highlighted in color and are underlined. Click on the hyperlink to go or open the corresponding location (for example: another section in the document, a website, or an e-mail address). |
| Boldface | Button labels are highlighted in boldface. For example, Start button. |

### 3. Purpose

This protocol generates ppmSeq™ libraries from genomic DNA (gDNA) using NEBNext UltraShear™, NEBNext® Ultra™ II End Repair/dA-Tailing, and NEBNext® Ultra™ II Ligation reagents. The resulting PCR-free libraries are compatible with the Ultima Genomics® UG 100™ Sequencer.

This protocol was developed based on the NEBNext UltraShear™ with NEBNext® Ultra™ II End Repair/dA-Tailing Module and NEBNext® Ultra™ II Ligation Module protocols with steps added or modified for sequencing compatibility on the UG 100™ Sequencing Platform.

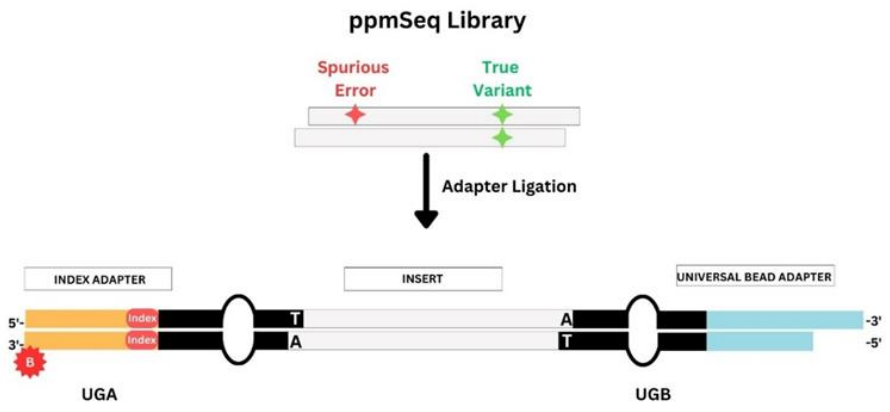

**Figure 1:** ppmSeq™ Library Workflow. This schematic illustrates the ppmSeq™ Library Workflow, designed for genomic DNA samples.

### 4. Support

Contact for support and assistance.

Mailing Address:

Ultima Genomics, Inc.
4209 Technology Drive
Fremont, CA 94538
USA

### 5. Workflow Overview

| ppmSeq™ Genomic DNA Library Preparation (16 Samples) | Approximate Time |
|---|---|
| **Enzymatic DNA Fragmentation** | |
| Add buffer and enzyme mixes to sample | **60 mins** |
| Run thermocycler | |
| **End Repair and dA-Tailing** | |
| Add buffer and enzyme mixes to sample | **75 mins** |
| Run thermocycler | |
| **Adapter Ligation** | |
| Add adapters to sample | |
| Add ligation master mix to sample | **80 mins** |
| Incubate at 25°C for 60 mins | |
| **Cleanup and Size-Selection** | |
| **SPRI Cleanup** | |
| Proceed with SPRI cleanup (0.79x). | |
| Transfer the supernatant to a new tube after the first SPRI cleanup stage | **90 mins** |
| **SPRI Size-Selection** | |
| Proceed with SPRI size-selection (0.65x/1x). | |
| Elute the size-selected library | |
| **Sample QC** | |
| Bioanalyzer, TapeStation, or Fragment Analyzer | **150 mins** |
| qPCR | |
| **Total Time** | **~7 hour 35 mins** |

Detailed breakdown of workflow steps, from input to quality control (QC), with estimated time and hands-on duration for 16 samples.

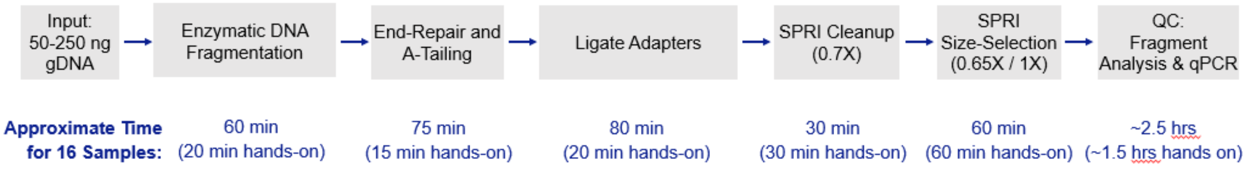

### 6. Materials Required

#### Reagents Supplied by Ultima Genomics

##### Indexed UG ppmSeq™ v3 Adapters

| Reagent Name | Part Number | Storage | Reaction Size |
|---|---|---|---|
| UG ppmSeq™ Adapter Plate v3 (96 Barcodes) | L21 | -20°C | 192 rxns per plate |
| UG ppmSeq™ Adapter v3 Tubes (16 Barcodes) | 500, 501, 502, 503, 504, 505, 506, 507, 508, 509, 510, 511, 512, 513, 514, 515 | -20°C | 8 rxns per tube |

For ordering inquiries related to Indexed UG ppmSeq™ v3 Adapters, please contact Ultima Genomics or reach out to your Field Application Scientist (FAS) from Ultima Genomics.

#### Customer Supplied Reagents

**NEBNext UltraShear™ (catalog #M7634)**

Included in the kit (information according to manufacturer):

| Contents | Storage |
|---|---|
| NEBNext UltraShear™ | -20°C |
| NEBNext UltraShear™ Reaction Buffer | |
| 500 mM DTT | |

**NEBNext® Ultra™ II End Repair/dA-Tailing Module (catalog #E7546)**

Included in the kit (information according to manufacturer):

| Contents | Storage |
|---|---|
| NEBNext®Ultra™ II End Prep Enzyme Mix | -20°C |
| NEBNext®Ultra™ II End Prep Reaction Buffer | |

**NEBNext® Ultra™ II Ligation Module (catalog #E7595)**

Included in the kit (information according to manufacturer):

| Contents | Storage |
|---|---|
| NEBNext®Ultra™ II Ligation Enhancer | -20°C |
| NEBNext®Ultra™ II Ligation Master Mix | |

Alternatively, NEBNext® Ultra™ II DNA PCR-free Library Prep Kit for Illumina (catalog #E7410) can be purchased in lieu of the NEBNext® Ultra™ II End Repair/dA-Tailing Module (catalog #E7546) and NEBNext® Ultra™ II Ligation Module (catalog #E7595).

**NEBNext® Library Quantification Kit for UG Libraries:** For ordering inquiries related to the NEBNext® Library Quantification Kit for UG Libraries, please contact your Field Application Scientist (FAS) from Ultima Genomics.

#### Additional Reagents

| Item | Source | Part Number |
|---|---|---|
| Agencourt™ AMPure™ xP Kit (or similar SPRI beads) | Beckman Coulter | A63880 |
| Agilent™ High Sensitivity DNA Kit or Agilent Genomic DNA ScreenTape | Agilent | 5067-4626 (or similar) |
| Ethanol, Absolute (200 Proof), Molecular Biology Grade | Fisher Scientific | BP2818100 |
| Buffer EB (10mM Tris-HCl, pH 8.5) | Qiagen | 19086 |
| 1 x TE (10 mM Tris pH 8.0, 1 mM EDTA) | Fisher Scientific | BP2473500 |

### Customer Supplied Consumables

| Item | Source | Part Number |
|---|---|---|
| Barrier Pipette Tips | Rainin | LTS Pipette Tips |
| 0.2 mL PCR Tubes or Plates | USA Scientific | 1402-2700, 1402-9100 or similar |
| Eppendorf LoBind™ Tubes, 1.5 mL | Eppendorf | 22431021 |

### Recommended Equipment

| Item | Source | Part Number |
|---|---|---|
| Pipettors 1–1000 µL | Rainin | 17014496, 30386739 |
| Magnetic Rack (for 0.2 mL tubes) | NEB | S1515S, or Thermo Scientific 492025 |
| Vortex Mixer, Vortex-Genie® 2 | USA Scientific | 7404-5600 |
| Microcentrifuge | USA Scientific | 2621-0016 |
| Agilent™ 2100 Bioanalyzer™ Instrument or Agilent™ 4150 TapeStation System | Agilent | G2939AA or G2992AA |
| Thermalcycler | BioRad | Heated lid, 0.2 µL tube (or similar) |
| Real-Time PCR (qPCR) Instrument | User-supplied | User-supplied |

### 7. Procedure

#### Fragment gDNA Using NEBNext UltraShear

1. For each sample, use 50-250 ng of purified genomic DNA (gDNA) diluted to a final volume of 26 µL in 1x TE (10 mM Tris pH 8.0, 1 mM EDTA).

**NOTE**

The fragmentation reaction is sensitive to the buffer composition, and poorly optimized fragmentation can result in low library yields. Only 1x TE (10 mM Tris pH 8.0, 1 mM EDTA) should be used to dilute the gDNA. If possible, the gDNA should be purified directly into 1x TE. If the purified gDNA is in another buffer and a significant volume will be added to the fragmentation reaction, it is advised that the gDNA be buffer exchanged into 1x TE. Furthermore, contaminants from certain gDNA extraction techniques can inhibit fragmentation. Performing a cleanup on the gDNA should rescue fragmentation.

2. Thaw NEBNext UltraShear™ Reaction Buffer completely, then vortex quickly to mix. Place on ice until use.
3. Vortex the NEBNext UltraShear enzyme for 5-10 seconds before use, and place on ice.
4. Program the thermocycler with the following conditions. Set the temperature of the heated lid to 75°C. Start the program to pre-chill the thermocycler.

| Step | Temp | Time |
|---|---|---|
| 1 | 4°C | HOLD |
| 2 | 45°C | 25 min |
| 3 | 65°C | 15 min |
| 4 | 4°C | Hold |

**NOTE**

If library yield is low, it may be necessary to optimize the fragmentation time at 45°C. Aim to maximize the amount of fragmented gDNA in the 250-270 bp size range.

5. Prepare the fragmentation reactions in 0.2 mL PCR tubes or plates on ice:

| Component | Volume Per Sample (µL) |
|---|---|
| gDNA sample | 26 |
| NEBNext UltraShear Reaction Buffer | 14 |
| NEBNext UltraShear | 4 |
| **Total Volume** | **44** |

6. Vortex the reactions for 5-10 seconds, and spin briefly in a microcentrifuge.
7. Place the samples in the pre-chilled thermocycler and proceed with steps 2-4 of the thermocycler program.

**SAFE STOPPING POINT!**

This is a safe stopping point. Libraries can be stored overnight at -20°C.

### End Repair and A-Tailing Using NEBNext Ultra II End Repair/dA-Tailing Module

**NOTE**
Only the NEBNext Ultra II End Prep Enzyme Mix and 500 mM DTT are used in this protocol. The NEBNext End Prep Reaction Buffer is not used.

1. Thaw the 500 mM DTT, then vortex to mix.
2. Prepare the End Repair and A-Tailing reactions on ice:

| Component | Volume Per Sample (µL) |
|---|---|
| Fragmented gDNA | 44 |
| 500 mM DTT | 2 |
| NEBNext Ultra II End Prep Enzyme Mix | 3 |
| **Total Volume** | **49** |

3. Set a pipette to 40 µL, then pipette the samples up and down at least 10 times to mix. Spin briefly in a microcentrifuge.

**NOTE**
Thorough mixing is critical for optimal performance. The presence of a small amount of bubbles will not compromise performance.

4. Place the samples in a thermocycler with the heated lid set to ≥75°C, and run the following program:

| Step | Temp | Time |
|---|---|---|
| 1 | 20°C | 30 min |
| 2 | 65°C | 30 min |
| 3 | 4°C | Hold |

5. Proceed immediately to Adapter Ligation.

### Adapter Ligation with NEBNext Ultra II Ligation Module

1. Thaw the UG ppmSeq™ Adapter Plate v3. Vortex to mix, and spin briefly in a microcentrifuge.
2. Mix the NEBNext Ultra II Ligation Master Mix by pipetting up and down several times before use.
3. Prepare Ligation reactions:

| Component | Volume Per Sample (µL) |
|---|---|
| End-Repaired and A-Tailed sample | 49 |
| Elution Buffer (10 mM Tris pH 8.0) | 7.25 |
| UG ppmSeq™ v3 Adapters (Indexed) | 6.25 |
| NEBNext Ultra II Ligation Master Mix | 30 |

| Component | Volume Per Sample (µL) |
|---|---|
| NEBNext Ligation Enhancer | 1 |
| **Total Volume** | **93.5** |

**NOTE**
The Ligation Master Mix and Ligation Enhancer can be mixed ahead of time. This premix is stable for at least 8 hours at 4°C. Do not add the adapters to the premix.

4. Set a pipette to 80 µL, then pipette the samples up and down at least 10 times to mix. Spin briefly in a microcentrifuge.
5. Place the samples in a thermocycler with the heated lid turned off, and run the following program:

| Step | Temp | Time |
|---|---|---|
| 1 | 20°C | 60 min |
| 2 | 4°C | Hold |

6. Proceed immediately to Library Clean-Up.

#### Library Clean-Up

1. Allow AMPure beads to warm to room temperature. Vortex to resuspend thoroughly.
2. Add 65.5 µL of AMPure beads (0.7 x final) to each ligation reaction.
3. Vortex the reactions for 10 seconds, then incubate for 5 minutes at room temperature.
4. Spin down, and place samples on magnet for 4 minutes.
5. Remove and discard supernatant.
6. Keeping samples on the magnet, add 200 µL of 80% ethanol. Incubate at room temperature for 30 seconds.
7. Remove and discard the ethanol supernatant.
8. Repeat wash: Keeping samples on the magnet, add 200 µL of 80% ethanol. Incubate at room temperature for 30 seconds.
9. Remove and discard the ethanol supernatant.
10. Spin down, place samples back on the magnet, and remove remaining ethanol. Air dry for 1 minute on magnet at room temperature.
11. Add 51 µL of Elution Buffer (10 mM Tris pH 8.0) to each sample.
12. Vortex 30 seconds, then incubate for 2 minutes at room temperature.
13. Spin down, and place samples on a magnet for 4 minutes.

**NOTE**
Do NOT discard supernatant!

14. Transfer the clear supernatant (50 µL) to a new tube, and discard beads.
15. Proceed to Size Selection or store overnight at -20°C.

**SAFE STOPPING POINT!**
This is a safe stopping point. Libraries can be stored overnight at -20°C.

#### Size Selection

1. Add 32.5 µL of AMPure beads to each sample (0.65 x final).
2. Vortex 30 seconds, then incubate samples for 5 minutes at room temperature.
3. Spin down, and place samples on a magnet for 4 minutes.

**NOTE**
Do NOT discard supernatant!

4. Transfer the supernatant (82.5 µL) to a new PCR tube, and discard beads.
5. Add 17.5 µL AMPure beads (1 x final) to the transferred supernatant.
6. Vortex 10 seconds, then incubate samples 5 minutes at room temperature.
7. Spin down, and place samples on a magnet for 4 minutes.
8. Remove and discard supernatant.
9. Keeping samples on the magnet, add 200 µL of 80% ethanol. Incubate at room temperature for 30 seconds.
10. Remove and discard the ethanol supernatant.
11. Repeat wash: Keeping the samples on the magnet, add 200 µL of 80% ethanol. Incubate at room temperature for 30 seconds.
12. Remove and discard the ethanol supernatant.
13. Spin down, and place samples back on the magnet.
14. Remove and discard the remaining ethanol. Air dry for 1 minute on the magnet at room temperature.
15. Add 63 µL Elution Buffer (10 mM Tris pH 8.0) to each sample.
16. Vortex for 30 seconds, then incubate samples for 2 minutes at room temperature.
17. Spin down, and place samples on a magnet for 2 minutes.

**NOTE**
Do NOT discard supernatant!

18. Transfer supernatant (63 µL) to a new tube, and discard beads.
19. Proceed to QC analysis (BioAnalyzer or TapeStation to determine mean fragment length, and qPCR to determine concentration)

**SAFE STOPPING POINT!**

This is a safe stopping point. Libraries can be stored for up to 1 week at 4°C or for 3 months at -20°C.

### 8. Library QC

1. Run 1μL of each library on a Bioanalyzer High Sensitivity Chip (or similar) to determine the mean library length in the range of 110-700 bp.

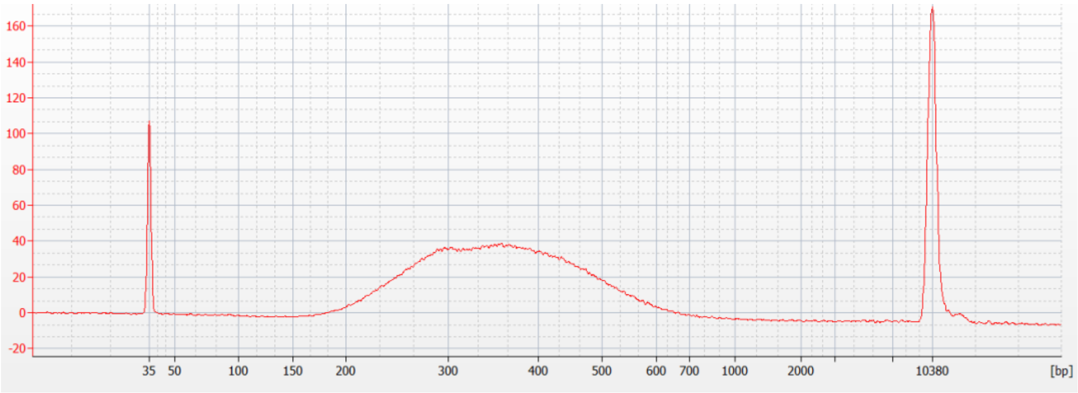

**Figure 2:** Example High Sensitivity Bioanalyzer trace of a ppmSeq™ library generated from 100 ng gDNA

2. Measure the library concentrations using the NEBNext® Library Quantification Kit for UG Libraries as described in the UG. Library Quantification Protocol (P00064).

### 9. Best Practices

The following recommendations are provided as a general guideline:

#### General Dilution Tips

1. Use freshly calibrated pipettes and certified free-from-DNAse/RNAse tips to prevent contamination and ensure accurate pipetting.
2. Minimize exposure to physical stress and light, especially for fluorescent dye-based methods, to prevent degradation and quenching.
3. Label all tubes clearly and keep a detailed log of dilutions to avoid sample mix-ups.
4. Work quickly but carefully to prevent evaporation, which can affect dilution factors, especially in small volumes.
5. Consider the impact of evaporation and adsorption in low-volume dilutions and use low-binding tubes and tips to minimize sample loss.

#### Dilution Guidelines and Factors for Library Preparation

Dilution Factors for Starting Samples:

DNA: For library preparation, an initial DNA concentration of 1-10 ng/µL is often required. If the starting DNA concentration is 50 ng/µL, a dilution of 1:5 or 1:10 may be appropriate to achieve this range.

General Dilution Guidelines:

1. Serial Dilutions: For samples with high concentrations, serial dilution is the preferred method to minimize error. A standard approach might be a 1:10 dilution followed by a 1:100 dilution, effectively resulting in a 1:1000 overall dilution.
2. Low-Volume Samples: When handling small amounts, it is advisable to avoid diluting less than 2 µL of the starting material or library into the final diluent volume, to prevent significant loss of volume during pipetting.

### 10. Safety and Regulatory

#### General

Using this product in a manner not specified in the user documentation may result in personal injury or damage to the instrument or device. Ensure that anyone using this product has received instructions in general safety practices for laboratories and the safety information provided in this document.

Wear appropriate personal protective equipment before working with reagents and operating the instrument.

#### Waste

Dispose waste in accordance with applicable regional, national, and local laws and regulations. Refer to kit SDS(s) for disposal considerations.

#### Safety Data Sheets (SDS)

Safety Data Sheets (SDS) for reagents described in this user guide can be found in the following link: https://regulatory.ultimagenomics.com/public

Contact for further questions regarding reagent SDS's.

#### Protocol Use

RUO – For Research Use Only. Not for use in diagnostic procedures.
